# Systems-level identification of key transcription factors in immune cell specification

**DOI:** 10.1101/2022.04.21.489000

**Authors:** Cong Liu, Kyla Omilusik, Clara Toma, Nadia S. Kurd, John T. Chang, Ananda W. Goldrath, Wei Wang

**Author notes:** These authors contributed equally to this work (Cong Liu, Kyla Omilusik).

## Abstract

Transcription factors (TFs) are crucial for regulating cell differentiation during the development of the immune system. However, the key TFs for orchestrating the specification of distinct immune cells are not fully understood. Here, we integrated the transcriptomic and epigenomic measurements in 73 mouse and 61 human primary cell types, respectively, that span the immune cell differentiation pathways. We constructed the cell-type-specific transcriptional regulatory network and assessed the global importance of TFs based on the Taiji framework, which is a method we have previously developed that can infer the global impact of TFs using integrated transcriptomic and epigenetic data. Integrative analysis across cell types revealed putative driver TFs in cell lineage-specific differentiation in both mouse and human systems. We have also identified TF combinations that play important roles in specific developmental stages. Furthermore, we validated the functions of predicted novel TFs in murine CD8+ T cell differentiation and showed the importance of Elf1 and Prdm9 in the effector versus memory T cell fate specification and Kdm2b and Tet3 in promoting differentiation of CD8+ tissue resident memory (Trm) cells, validating the approach. Thus, we have developed a bioinformatic approach that provides a global picture of the regulatory mechanisms that govern cellular differentiation in the immune system and aids the discovery of novel mechanisms in cell fate decisions.

## Introduction

The immune system protects the human body by eliminating invading pathogens as well as tumors[1,2]. Considerable effort has been devoted to developing preventative and therapeutic treatments based on exploitation of a diverse set of immune cell types. Establishing immune cell-based clinical treatments relies on research advances that reveal the fundamental mechanisms regulating the immune system, such as those that direct the differentiation from progenitor to mature immune cells and coordinate their response to the presence of various antigens. Recent technological advances have made it possible to study the regulatory mechanisms at the systems level and genomic scale[3]. However, the noise in the genomic measurements and the complexity of integrating multiple omics assays pose a great challenge of fully digesting the information and extracting the most informative signals from the extensive data.

A critical step towards achieving the goal of studying the regulatory mechanism is to uncover the key regulators that direct the specification of immune cell states. Transcription factors (TFs) are essential regulators of cell fates[4] and play pivotal roles in immune cell development[5]. Identification of critical TFs that drive immune cell differentiation can also provide key mechanistic insights into immune cell functions. However, a complete catalog of critical TFs in each immune cell population or in each lineage has not been established. Therefore, dissection of the genetic networks that promote immune cell differentiation and function and characterization of the genome-wide influences of the key regulators would provide novel insights into cell specification and potential strategies to impact immune function.

The global influence of a cellular regulator is conveyed through its regulatory effect on target genes, which is subsequently propagated over the genetic network. A given regulator’s activity is not only affected by its own expression level but also post-translational modifications, the presence of collaborative co-factors and its targets’ accessibility. Therefore, the expression level of a regulator such as a TF is not always correlated with its activity[6]. In light of this, many methods have been proposed to infer the activity of regulators using statistical or machine learning approaches. For instance, Schacht et al.[6] developed a statistical model to estimate the regulatory activity of TFs using their cumulative effects on their target genes. Arrieta-Ortiz et al.[7] used a linear model to infer the TF activity (TFA) by predicting target genes’ expression levels. SCENIC[8] developed by Aibar et al. constructs the genetic network by predicting each gene’s expression using the TF’s expression levels and finding the most predictive TFs as the key regulators. Maslova et al. introduced AI-TAC[9] to predict ATAC-seq signals and identified the most enriched motifs when evaluating the TF importance. Although these methods were able to predict the local activity of a TF, i.e. the expression level of their direct target genes, measuring the system-wide influence of a given TF was not their focus. As genes rarely function alone but usually cross-talk with each other to form complex regulatory logic, we argue that the global influence of a regulator should, in principle, better predict the cell state change upon perturbing the regulator than its local influence alone[10,11].

Recently, we have developed a method, called Taiji, that can successfully infer the global impact of TFs in the genetic network constructed from integrating epigenomic and gene expression data[12–14]. Focusing on the global rather than the local importance of transcriptional regulators makes Taiji robust and suitable for integrating multiomics data from noisy genomic measurements. In fact, Taiji clearly outperforms the motif enrichment analysis and the TFA approach[12–14]. In addition, several TFs predicted to be critical in regulating the differentiation of CD8^+^ T cells following infection were validated experimentally and demonstrated to have previously unappreciated roles in cell fate specification[13,14].

Here, we performed a comprehensive analysis on the common regulatory code in the mouse and human immune systems using Taiji. We identified key TFs critical for programming the differentiation of nine cell lineages that span diverse cell types of the immune system, thus providing the first comprehensive catalog of key regulators for a variety of cell lineages during immune system development. In addition, we uncovered TF combinations that activate in a spatiotemporal manner, behaving like transcriptional waves to orchestrate the developmental progress and cell lineage specification. We show the great predictive performance of Taiji by validating the functions of four novel TFs in murine CD8^+^ T cell differentiation and memory formation.

## Materials and Methods

### Dataset acquisition

The mouse dataset was downloaded from Gene Expression Omnibus (GEO) (GSE100738 and GSE109125 corresponding to ATAC-seq and RNA-seq, respectively[15]). We matched the ATAC-seq and RNA-seq by sample name and 115 samples in 73 cell types were used as input to Taiji. The cell types span nine cell lineages: stem cells, dendritic cells (DC), macrophages/neutrophils (MF/GN), monocytes (Mo), B cells, innate lymphoid cells (ILC), αβ T cells (abT), γδ T cells (gdT), and activated T cells (Act_T).

The human datasets analyzed in this study were retrieved from GEO (GSE118189 and GSE118165, GSE74912 and GSE74246)[16,17]. In total, 226 samples distributed in 61 immune cell types were chosen as input. The cell types span nine cell lineages: B, CD4T, CD8T, γδT, monocytes, dendritic cells, natural killer (NK), and stem cells. All of the sample names and division methods were consistent with the original paper.

### TF regulatory networks construction and visualization

Taiji v1.1.0 with default parameters was used for the integrative analysis of RNA-seq and ATAC-seq data (https://github.com/Taiji-pipeline/Taiji). The motif file was downloaded directly from the CIS-BP database. There were 815 mouse motifs. The majority of these proteins are TFs.

The network shown in Fig. 3c is a subset of the whole network for PBX4 in T8.IEL.LCMV.d7.SI. We first removed the edges with weight less than 1 and then retained 500 out of 1437 regulatees via random sampling while keeping the ratio between non-TF nodes and TF-nodes constant. Next, we constructed the regulatory network between TF-nodes and non-TF nodes within 500 regulatees and thus expanded the network to 4213 edges. For better visualization we further downsampled 25% of the total edges and used it as input. Cytoscape v3.8.0 was used for network visualization. The size of the node is proportional to its PageRank score.

### TF regulatory networks weighting scheme

As described in the original Taiji paper[12], a personalized PageRank algorithm is applied to calculate the ranking scores for TFs. We first initialized the edge weights and node weights in the network. The node weight was calculated as e^zi^, where z_i_ is the gene’s relative expression level in cell type i, which is computed by applying the z score <u>t</u>ransformation to its absolute expression levels. The edge weight is determined by e_ij_ = 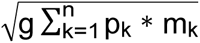, where p is the peak intensity, represented by the p-value of the ATAC-seq peak at the predicted TF binding site, rescaled to [0, 1] by a sigmoid function; m is the motif binding affinity, represented by the p-value of the motif binding score, rescaled to [0, 1] by a sigmoid function; g is the TF expression value; n is the number of binding sites linked to gene j.

If the TFs in the same protein family share the same motifs, their PageRanks scores are distinguished by their own expression levels because their motifs and the target genes are the same. If a motif is weak, the PageScore of the TF is decided by whether these motifs occur in the open chromatin regions (measured by the peak intensity of the ATAC-seq data), the TF expression and its target expression levels. Because we also compare TFs between different cell types to identify cell-type/lineage specific regulators. The relative difference between the PageRank scores of TFs also helps to uncover important TFs with weak motifs.

### Identification of putative driver TFs in cell lineage specification

To identify lineage-specific TFs, we divided the samples into two groups: target group and background group. Target group included all the samples belonging to the cell type/lineage of interest and the background group comprised the remaining samples (Table S1). We then performed the normality test using Shapiro-Wilk’s method to determine whether the two groups were normally distributed and we found that the PageRank scores of most samples (90%) follow log-normal distribution. Based on log-normality assumption, an unpaired t-test was used to calculate the P-value. A P-value cutoff of 0.001 and log_2_ fold change cutoff of 1 were used for calling lineage-specific TFs (see the full result in Fig. S2).

### Identification of memory cell-specific and tissue-resident-specific TFs

To identify memory cell-specific TFs, we grouped three memory cells, i.e. B.mem.Sp, T8.Tcm.LCMV.d180.Sp, T8.Tem.LCMV.d180.Sp, into target group while leaving other samples except for T8.MP.LCMV.d7.Sp as background, because T8.MP.LCMV.d7.Sp is a memory precursor cell. The normality test showed that most samples were log-normal distributed so that we performed unpaired t-test based on log-normality assumption. A P-value cutoff of 0.1 was used to identify the memory cell-specific TFs. The list was filtered by removing the digestion-related genes. We identified 30 memory cell-specific TFs.

To identify tissue-resident-specific TFs, we grouped six lymphocytes resident in small intestine (ILC3.NKp46+.SI, ILC3.NKp46-CCR6-.SI, ILC3.CCR+.SI, ILC2.SI, Treg.4.FP3+.Nrplo.Co, T8.IEL.LCMV.d7.SI) as target group and eight similar cells in circulation as background (T8.Tcm.LCMV.d180.Sp, T8.Tem.LCMV.d180.Sp, T8.MP.LCMV.d7.Sp, T8.TE.LCMV.d7.Sp, Treg.4.25hi.Sp, NK.27+11b-.Sp, NK.27+11b+.Sp, NK.27-11b+.Sp). Based on the log-normality assumption, we performed the unpaired t-test and used double cut-off (P-value <0.1 and log2 fold change > 0.5) for calling the putative driver TFs.The list was filtered by removing the digestion-related genes. A list of 54 tissue-residency-specific TFs were thus identified.

### GO and pathway enrichment analysis

R package clusterProfiler 4.0 (https://bioconductor.org/packages/release/bioc/html/clusterProfiler.html) was used to perform the enrichment analysis of the identified TFs, in which KEGG Pathway (http://www.genome.jp/kegg/catalog/org_list.html) and DAVID (via RDAVIDWebService https://bioconductor.org/about/removed-packages/). A cutoff of P-value < 0.1 was used to select the significantly enriched pathways.

### Mouse and Cell lines

All mouse strains were bred and housed in specific pathogen–free conditions in accordance with the Institutional Animal Care and Use Guidelines of the University of California San Diego. C57Bl/6J, P14 (with transgenic expression of H-2D^b^-restricted TCR specific for LCMV glycoprotein gp_33_), CD45.1^+^ and CD45.1.2^+^ congenic mice were bred in house. Both male and female mice were used throughout the study, with sex- and age-matched T cell donors and recipients.

### Infections Studies

shRNAmirs targeting mouse *Elf1*, *Prdm9*, *Kdm2b*, *Tet3* or *Cd19* in a pLMPd-Amt vector were used to generate retroviral supernatant as previously described[13]. Naive P14 T cells from the spleen and LN were enriched via negative selection using MACS magnetic beads following the manufacturer’s protocol (Miltenyi Biotec) then activated at 2×10^6^ cells/well in six-well plates coated with 100μg/mL goat anti-hamster IgG (H+L, Thermoscientific), 1μg/mL anti-CD3 (145–2C11), and 1μg/mL anti-CD28 (37.51) (eBioscience) for 18 h. Retroviral supernatant supplemented with of 50μM beta-mercaptoethanol and 8μg/mL polybrene (Millipore) replaced the culture media and cells were centrifuged for 60 min at 2,000 rpm, 37 °C. After 24 h, P14 T cells transduced with gene targeting or *Cd19* control shRNAmir encoding-retroviruses were mixed at 1:1 ratio and a total of 5×10^5^ cells were transferred to congenically distinct recipient mice that were infected with 2×10^5^ plaque-forming unit (PFU) LCMV-Armstrong i.p. A portion of the P14 CD8^+^ T cells transduced with gene targeting or *Cd19* control shRNAmir encoding-retroviruses were cultured *in vitro* for another 24h in media supplemented with 50 U/ml of human IL-2 and sorted CD8a^+^Ametrine^+^ cells were evaluated by qPCR for knockdown efficiency using the following primer pairs: *Elf1* F:CCAAGTATTAGGACTATACAGG, *Elf1* R:GGCTATAACCGTTGTGAGTG; *Prdm9* F:ATATGGAATGGAATCATCGC, *Prdm9* R:GTGCTG GGAAAGGTTGTT; *Kdm2b* F:AGCAGACAGAAGCCACCAAT, *Kdm2b* R:AGGTGCCTCCAAAG TCAATG; *Tet3* F:TGCGATTGTGTCGAACAAATAGT, T*et3* R:TCCATACCGATCCTC CATGAG.

### Preparation of Single Cell Suspensions

Single-cell suspensions were prepared from spleen by mechanical disruption. For small intestine preparations, Peyer’s patches were excised, and luminal contents were removed. The tissue was cut longitudinally then into 1cm pieces then incubated in 10% HBSS/HEPES bicarbonate solution containing 15.4mg/100mL of dithioerythritol (EMD Millipore) on a stir plate with rotating stir bar at 37°C for 30 min to extract intraepithelial lymphocytes (IEL). Gut pieces were further treated with 100U/ml type I collagenase (Worthington Biochemical) in RPMI-1640 containing 5% bovine growth serum, 2 mM MgCl_2_ and 2 mM CaCl_2_ at 37°C for 30 min. Salivary gland and kidney were cut with scissors into fine pieces then incubated while shaking with 100U/ml type I collagenase at 37°C for 30 min. Lymphocytes from all tissues but spleen were purified on a 44%/67% Percoll density gradient.

### Flow Cytometry and Cell Sorting

The following antibodies (from eBioscience and Biolegend) were used for surface staining: CD8a (53-6.7), CD8b (eBioH35-17.2), CD45.1 (A20-1.7), CD45.2 (104), CD62L (MEL-14), CD69 (H1.2F3), CD103 (2E7,), CD127 (A7R34), KLRG1 (2F1). Cells were incubated for 30 min at 4°C in PBS supplemented with 2% bovine growth serum and 0.1% sodium azide. To identify tissue-resident CD8^+^ T cells, 3 mg of CD8a (53–6.7) conjugated to APC eFlour780 was injected into mice three minutes prior to sacrifice and organ excision. CD8b^+^ cells without CD8a labeling were considered to be localized within non-lymphoid tissues. Stained cells were analyzed using LSRFortessa or LSRFortessa X-20 (BD) and FlowJo software (TreeStar). Cell sorting was performed on BD FACSAria Fusion instruments.

### Quantification and Statistical Analysis

Statistical analysis was performed using GraphPad Prism software. A one-way paired ANOVA or two-tailed ratio paired t-test was used for comparisons between groups as indicated in figure legends. P values of <0.05 were considered significant.

## Results

Using assay of transposase-accessible chromatin with high-throughput sequencing (ATAC-seq) and gene expression (RNA-seq) data for 73 mouse immune cell populations generated by the Immunological Genome Project[15], representing 9 cell types including stem cells, dendritic cells (DC), macrophages/neutrophils (MF/GN), monocytes (Mo), B cells, innate lymphoid cells (ILC), αβ T cells (abT), γδ T cells (gdT), and activated T cells (Act_T), we performed an integrated analysis using the Taiji pipeline[12]. Briefly, Taiji integrates gene expression and epigenetic modification data to build gene regulatory networks. Taiji first scans putative TF binding sites in each open chromatin region using motifs documented in the CIS-BP database[18], to identify active regulatory elements, including active promoters or enhancers. These TFs are then linked to their target genes as predicted by EpiTensor[19]. All of the regulatory interactions are assembled into a genetic network. Lastly, the personalized PageRank algorithm is used to assess the global influences of the TFs[20]. In the network, the node weights were determined by the z scores of gene expression levels, allocating higher ranks to the TFs that regulate more differentially expressed genes. The edge weights were set to be proportional to TFs’ expression levels, the binding site peak intensity, and the motif binding affinity, thus representing the regulatory strength (Fig. 1). The details of the weighting scheme are available in “Materials and Methods” section. The superior performance of Taiji compared to other methods to identify key regulators have been confirmed using simulated data, literature evidence and experimental validations[12,14]. We constructed a network in each mouse cell type. The average number of nodes and edges of the networks were 19,340 and 773,653, respectively, including 815 (4.21%) TF nodes. On average, each TF was predicted to regulate 3996 genes, and each gene was predicted to regulate 88 TFs (Fig. S1).

**Fig. 1.**
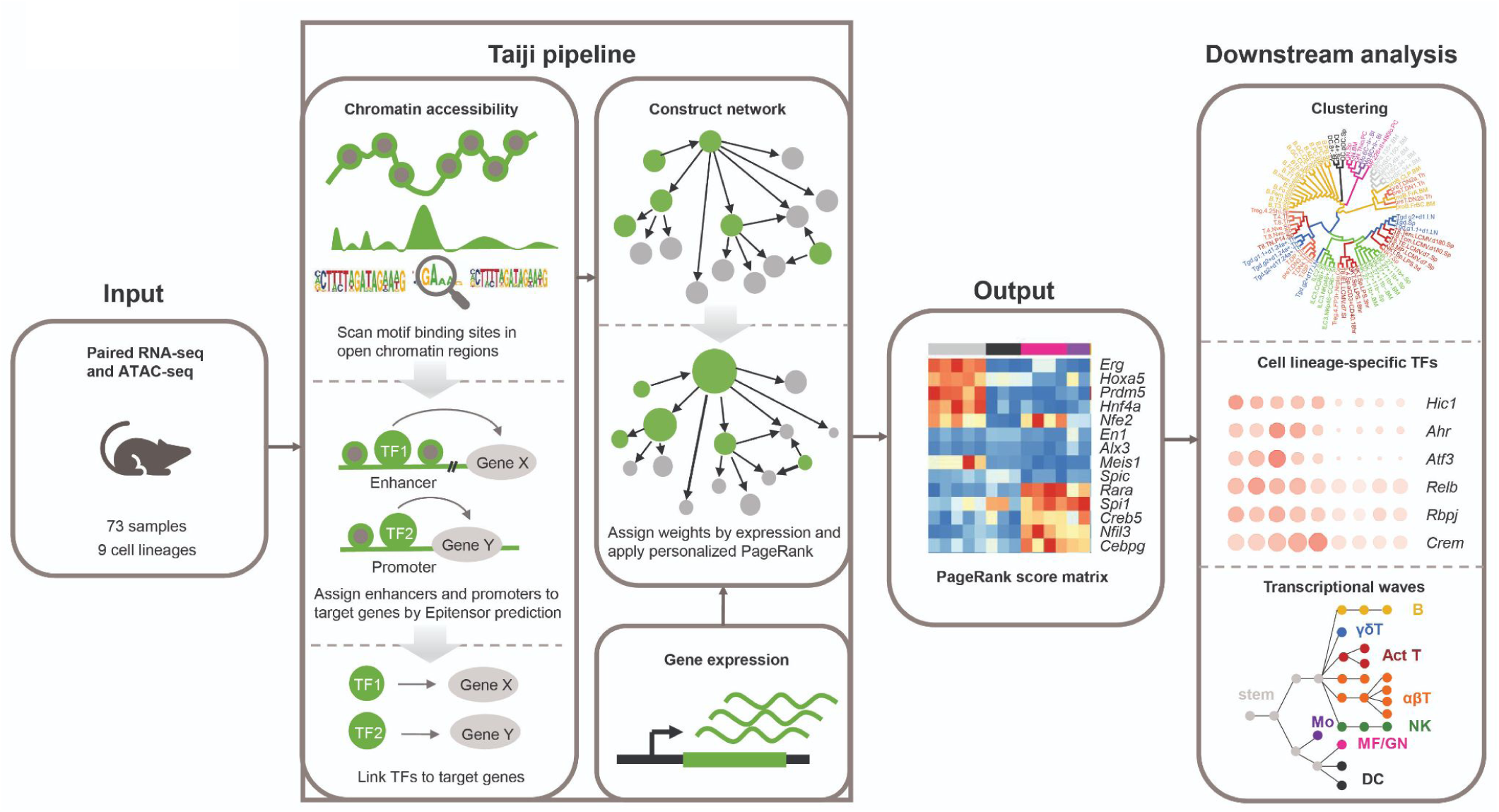
The workflow of study and Taiji pipeline. Matched RNA-seq and ATAC-seq data were used as input for the Taiji framework to construct a regulatory network and generate PageRank scores matrix as output (column colors code for different cell lineages, see Fig. 2 for the color annotation). Downstream analyses included clustering samples to distinct cell lineages, identification of cell lineage-specific TFs, and construction of temporal transcriptional waves.

### PageRank score is a great indicator of cell specificity

To characterize the potential global influences of all the TFs across different cell types, we removed TFs that do not present large variations (coefficients of variance (CVs) less than 1) in their PageRank scores, which gave us a list of 167 most variable TFs. We plotted the PageRank score matrix with the columns sorted by the nine cell lineages (Fig. 2). For visualization, we only showed 67 TFs among all the most variable TFs. These 67 TFs have CVs larger than 1 and less than 1.3. We found that different cell lineages show quite distinct TF regulatory patterns, illustrating that transcriptional regulation takes place in a cell-lineage-specific fashion. The row-wise comparison demonstrates that some TFs are specific to one particular cell lineage. For example, *Bcl11a* is essential for the survival of early B cells[21,22] and *Irf4* has been shown as a key transcription factor that directs the development and maturation of B cells including pre-B differentiation and marginal zone B cell development.[23–25]. Both *Bcl11a* and *Icf4* have exclusively higher PageRank scores in B cell lineage compared to the other immune lineages. Another example is that both *Tcf7* and *Lef1* have known functions in T cell development and differentiation[26–28] and they display higher PageRank scores across T cells compared to the average.

**Fig. 2.**
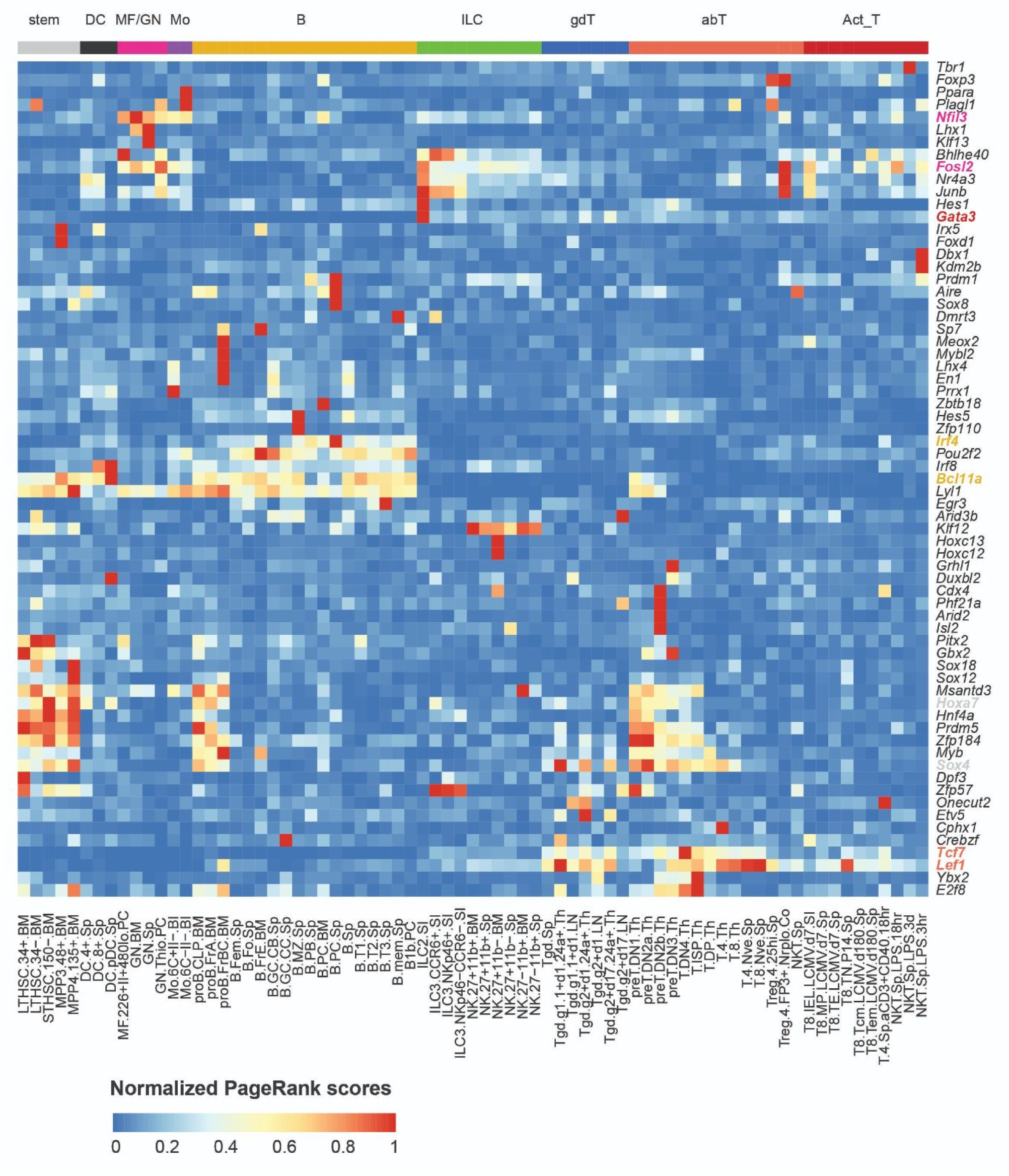
PageRank score is a great indicator of cell specificity. PageRank scores heatmap of 67 most variable TFs (1 < coefficients of variance (CVs) < 1.3) across 73 cell types. TFs in rows (z-normalized), cell types in columns, and color of the cell in the matrix indicates the normalized PageRank scores with red displaying high scores. Selected gene names are colored based on their cell specificity. abT, αβ T cells; Act_T, activated T cells; B, B cells; DC, dendritic cells; gdT, γδ T cells; ILC, innate lymphoid cells; MF, macrophages; GN, neutrophils; Mo, monocytes; stem, stem cells.

### Identification of putative driver TFs in cell lineage differentiation

We then identified 48 constitutively active TFs whose average PageRank scores across all the cell types were ranked top 10% with CVs<0.5, indicating their predictive activities across all lineages (Fig. 3a). Similar to their transcriptional activities, the expression levels of these TFs remained relatively high in immune cells from different tissues and at different stages of development (Fig. S1d, the circle size represents the expression levels while the color represents the PageRank scores). Functional analysis revealed that these TFs are enriched in house-keeping functions, including “biological regulation”, “metabolic process”, and “cellular process”, demonstrating the general roles of active TFs in basic cellular functions across all cell types. Notably, more than one third of these active TFs are related to the immune system processes, which is not surprising.

**Fig. 3.**
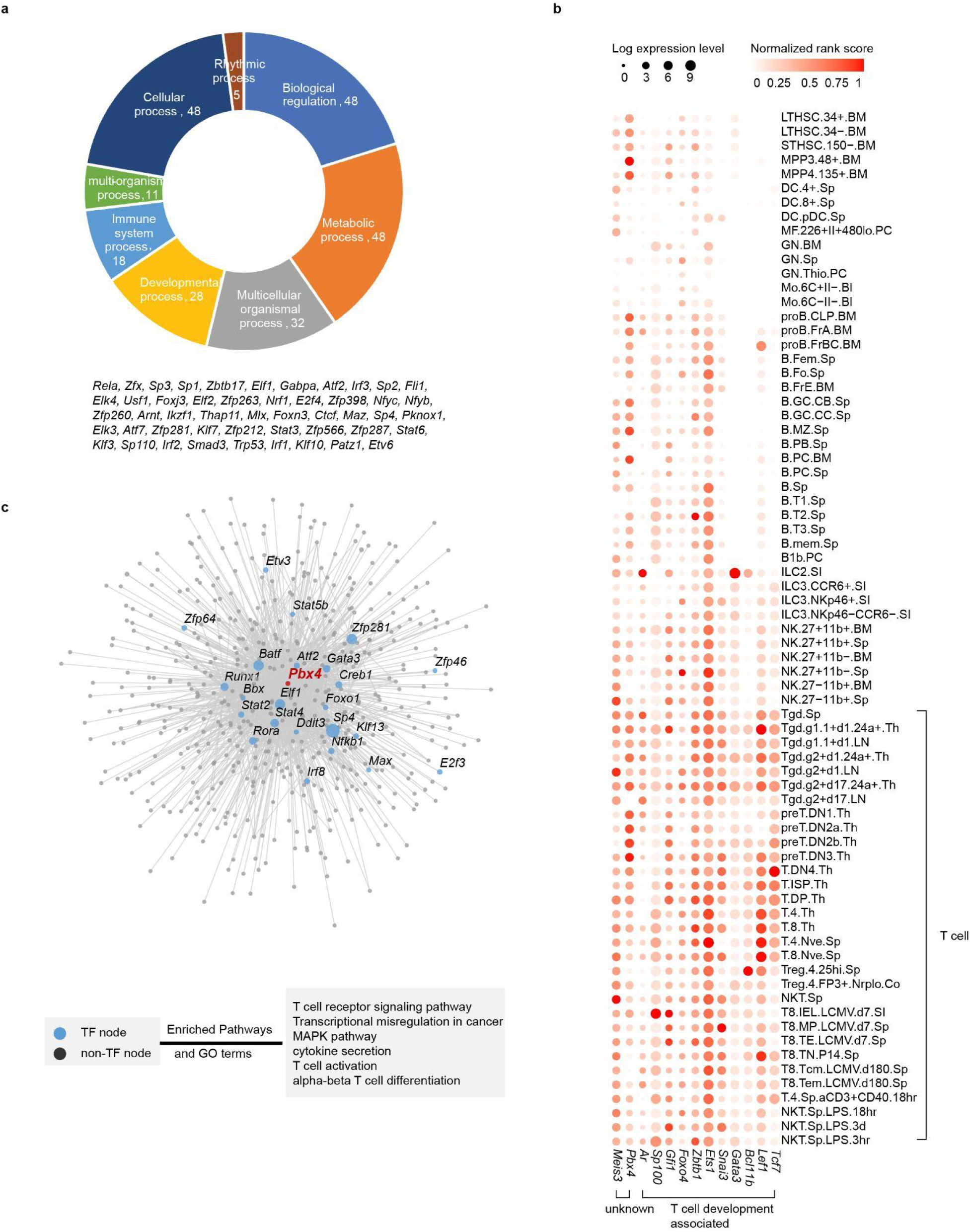
Identification of putative driver TFs in cell lineage specification during mouse immune system development. **(a)** Functional classification of 48 constitutively active TFs reveals their functions in essential biological processes. **(b)** Identified putative driver TFs in T cell lineage, including eleven TFs known to be related to T cell development, and two TFs with unknown functions. Circle size represents the logarithm of gene expression and the color represents the normalized PageRank score. **(c)** The network for *Pbx4*, a previously unknown putative driver TF in T cell lineage, and its regulatees in the T8.IEL.LCMV.d7.SI. Each node represents one regulatee of *Pbx4* with TF highlighted in blue. Larger node size represents a higher PageRank score in the network. The bottom box shows the four enriched Gene Ontology (GO) terms and KEGG pathway for *Pbx4*’s regulatees.

Furthermore, we identified potential important TFs in 73 cell types from 9 cell lineages by comparing their PageRank scores between a specific lineage and the background (see “Methods” for details). To our knowledge, this is the first systematic mapping of the complex transcriptional regulation that occurs during mouse immune system development (Fig. S3). The identified putative driver TFs include many well-known key regulators. For instance, 11 of 13 identified T cell-specific TFs have been previously reported to play a pivotal role during the T cell development and differentiation in the mouse immune system[29–31] (Fig. 3b), such as *Lef1* and *Tcf7* that are critical for CD8^+^ T cell memory formation and CD4^+^ T cell differentiation[32–36], and *Bcl11b*, an essential TF for the development of αβ T cells and most γδ T cells[37].

To systematically validate the predictions of these analyses, we performed literature search in 5 lymphoid hematopoietic lineages that have been extensively studied, including the αβT cells, Act_T cells, B cells, γδT cells, and ILC. Forty-two of 50 (84%) TFs identified by our method are shown to be associated with regulation of development or normal functions of these cell lineages (Table S2). For example, *Tcf4* is a well-known transcriptional activator in B cell differentiation and regulates the expression of genes critical for germinal center (GC) B cell development[38–41].

The lineage specificity was decided by assessing whether the TF PageRank scores are statistically significantly higher in one lineage than the other lineages, i.e. the p-value smaller than 0.001 and the log2 fold change larger than 1. It is possible that some TFs are important in several cell lineages. Take *Bcl11a* as an example, it exhibits high PageRank scores in several cell lineages: stem, DC cells and B cells, and its role in all these three lineages have been previously described[42–46]. We found that several putative driver TFs are highly conserved in three T cell subsets. *Lef1*, *Gata3*, and *Bcl11b* were ranked top 10 in all 3 T cell sub-lineages, demonstrating that interplay and co-expression of these regulators are essential for the development and differentiation of several T cell subsets including CD4^+^ and CD8^+^ T cells.

We have also identified TFs with previously undefined functions in immune cell differentiation. For example, *Pbx4*, identified as a key regulator for T cell development, has not yet been studied. We predicted that *Pbx4* regulates a number of high-rank TFs, including several aforementioned constitutively active TFs, i.e., *Sp4*, *Elf1*, and *Zfp281* (Fig. 3c). The functional enrichment analysis of *Pbx4*’s regulatees suggests that they participate in T cell receptor signaling pathway, T cell activation, and αβ T cell differentiation (Fig. 3c).

During development, hematopoietic stem cells (HSCs) differentiate into two different types of immune cells—myeloid and lymphoid lineages. Myeloid cells include Mo, DC, neutrophil, GN, and MF while lymphocytes include ILC, T and B lineages. T cell lineage can be further divided into abT, gdT, and Act_T. To identify TFs that are specific to each cell lineage, we first grouped and compared samples originated from myeloid lineage against samples from lymphoid lineage. The student t tests with a P value cutoff of 0.005 and log_2_ fold change cutoff of 0.5 were used to identify TFs whose ranks change significantly in myeloid lineage compared with lymphoid lineage and vice versa, thus obtaining a list of myeloid-specific TFs and lymphoid-specific TFs (Fig. 4). The functional relevance of the predicted specific TFs is supported by previous studies. For example, we have identified *Hhex* and *Spi1* as putative specific regulators for myeloid cells, which are known to be associated with myeloid leukocyte differentiation[47,48]. The enriched Gene Ontology (GO) terms showed that the identified lymphoid-specific TFs are involved in T cell receptor V(D)J recombination, B cell lineage commitment, and cellular response to interleukin-4.

**Fig. 4.**
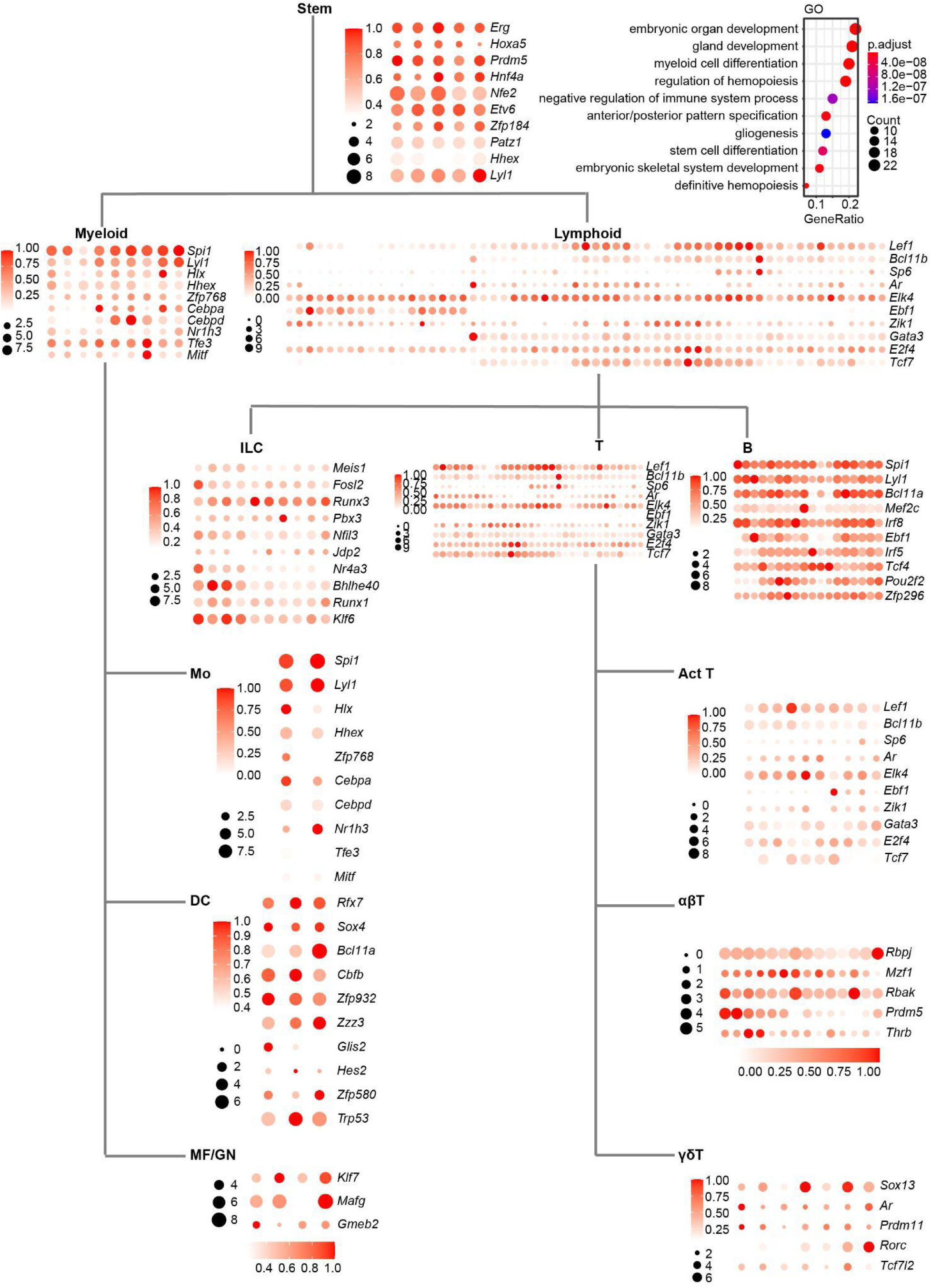
Cell lineage-specific putative driver TFs in the mouse immune system. A P-value of 0.005 and log2 fold change of 0.5 were used to call the putative driver TFs in each cell lineage. The bubble plots show the top 10 putative driver TFs because of the limited space. The GO term enrichment analysis for stem cells is shown as an example.

To analyze the finer cell lineage specificity of the predicted putative driver TFs, we further compared each sub-cell lineage with other cell lineages originated from the same progenitors and used the student t tests with a P value cutoff of 0.005 and log_2_ fold change cutoff of 0.5 to identify cell lineage-specific putative driver TFs. In total, 241 cell lineage-specific putative driver TFs were found and the top 10 TFs are shown in Fig. 4. GO terms and pathway enrichment analysis is performed for each group of cell lineage-specific putative driver TFs (Table. S4). Many well-known cell lineage-specific markers appeared in the list, e.g., *Fosl2*[49] and *Runx3*[50,51] for NK cells and *Bcl11a*[46] for DC. In addition, a number of novel TFs were also found from our analysis, such as *Zfp296* for B cell, *Jdp2* for NK cell and *Ar* for Act T cells. Taken together, these results provide a comprehensive map for the future mechanistic study of mouse immune system development.

### Transcriptional waves during immune cell development

In addition to the tissue specificity, we have also analyzed the dynamic activity of TFs during immune cell development. To identify the TFs which show similar temporal activity patterns, we clustered the TFs based on the normalized PageRank scores across samples. First of all, we performed the principal component analysis (PCA) for dimension reduction of the TF score matrix. We retained the first 30 principal components for further clustering analysis, which explained 75% variance (Fig. S3a). We used the k-means algorithm for clustering analysis. To find the optimal number of clusters and similarity metric, we performed the Silhouette analysis to evaluate the clustering quality using five distance metrics: Euclidean distance, Manhattan distance, Kendall correlation, Pearson correlation, and Spearman correlation (Fig. S3b). Pearson correlation was the most appropriate distance metric since the average Silhouette width was the highest among the five distance metrics. Based on these analyses, we identified 40 distinct dynamic patterns of TF activity during immune cell development (Fig. S3).

The 40 clusters represent transcriptional waves that orchestrate tissue differentiation. Some TFs are predicted to be active throughout many developmental stages, represented by cluster C21 and C29 (Fig. 5a, b). In both clusters, TFs exhibit relatively high PageRank scores and show active activity across almost all stages. Example TFs include constitutively active genes identified earlier, such as *Sp1*[52], *Zfx*[53], and *E2f4*[54], which are essential for immune system development and maintenance of normal functions of various immune cells.

**Fig. 5.**
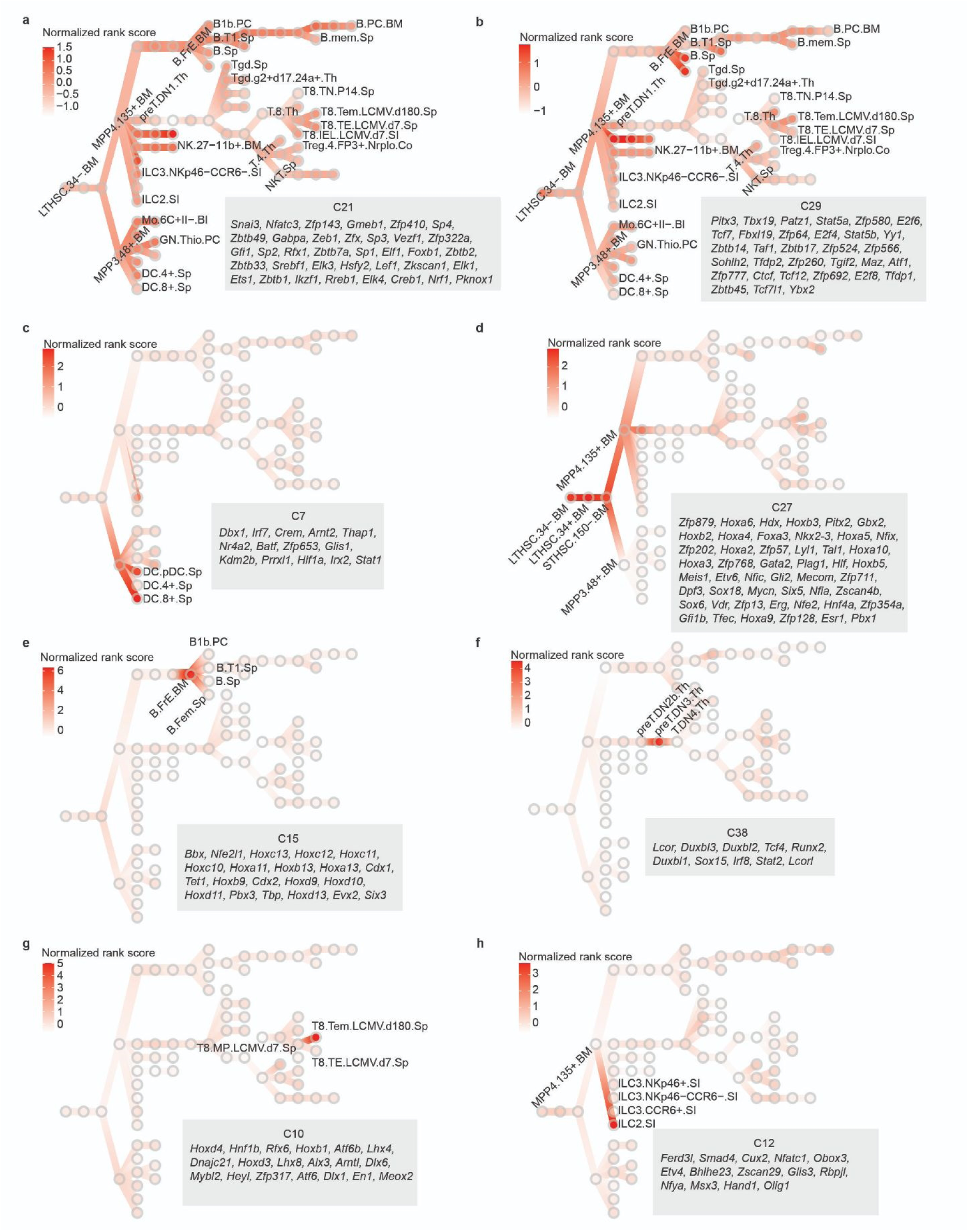
Transcriptional waves regulate the development of the mouse immune system. **(a,b)** Two transcriptional waves C21 and C29, which are active in all developmental stages. **(c,d)** Two cell lineage-specific transcriptional waves. C7 is active in DC lineage and C27 is active in stem cells. **(e-h)** Four examples of stage-specific waves in immune cell development. C15 is active in B.FrE.BM; C38 is active in preT.DN3.Th; C10 is active in T8.Tem.LCMV.d180.Sp; C12 is active in ILC2.SI. Circles in each panel represent specific cells in the developmental stages. Color indicates normalized PageRank scores with red displaying high values. TF members of the cluster are shown on the bottom right of the panel.

We also found clusters that include genes responsible for the differentiation of specific cell lineages. For example, C7 (Fig. 5c) is particularly active in DC samples, suggesting its critical role in DC cell differentiation. Many TFs that are crucial for DC cell development were recovered in C7, such as *Stat1*[55], *Irf7*[56], and *Batf*[57]. Another instance of cell lineage-specific clusters is C27 (Fig. 5d), C27 is exclusively active in stem cells, particularly in 3 HSCs (LTHSC.34-.BM, LTHSC.34+.BM, STHSC.150-.BM,). TFs in this cluster are associated with functions specific to early development of HSC including ten *Hox*[58] family TFs and two *Sox*[59] family TFs.

In addition to cell lineage–specific transcriptional waves, there are also developmental stage-specific transcriptional programs that appear to drive the differentiation of individual stages. We highlighted 4 clusters as examples, C15, C38, C10, and C12, which are potentially responsible for the B.FrE.BM, preT.DN3.Th, T8.Tem.LCMV.d180.Sp, and ILC2.SI development, respectively (Fig. 5e-h). TFs in these clusters exhibit transient transcriptional spikes, likely related to their stage-specific functions, thus activating at different stages in immune system development. C38 includes three *Duxbl* family members (*Duxbl1*, *Duxbl2*, and *Duxbl3*), which mediate elimination of pre-T cells that fail β-selection[60]. Several *Hox* family TFs (*Hoxd4*, *Hoxb1*, *Hoxd3*) in C10 show significantly higher Pagerank score in CD8^+^ effector memory T cell (T_EM_) than that in others, suggesting their putative role in regulating T_EM_ development. Although previous evidence shows that *Hoxb4* overexpression can increase the central memory population in CD4^+^ T cell[61], the mechanism of *Hox* family TFs in CD8^+^ T cell has not been thoroughly explored. As the mechanism of immune system development remains to be completely defined, our discovery, therefore, provides an invaluable resource for future studies.

### Taiji identifies key TFs important for differentiation of effector and memory CD8^+^ T cell populations

Memory cells that mount a rapid and protective response upon reinfection are the fundamental basis for vaccines, long-term immunity and health[62]. To examine key regulators of memory formation, we further investigated TFs that are important for directing differentiation of short-lived effector versus long-lived memory cells. We identified putative driver TFs in 3 memory cell populations (B.mem.Sp, T8.Tcm.LCMV.d180.Sp, T8.Tem.LCMV.d180.Sp) with all other cells as background following the similar procedure to the identification of cell lineage-specific TFs (see Methods). We then performed literature search on the top 10 putative driver TFs (Table S3) and found that half of them are well-known regulators of the differentiation and maintenance of normal functions of memory T cells and B cells, such as *Foxo1*[63–70], *Ikzf3*[71,72], and *Ets1*[73], while others remain unknown and warrant a more thorough investigation, such as *Cphx1*, *Pou4f1*, and *Osr2*.

We selected 2 TFs predicted by Taiji to regulate CD8^+^ T cell differentiation with previously unknown functions in CD8^+^ T cell differentiation and memory formation – *Elf1* and *Prdm9* - to further validate our analysis. Congenically distinct P14 T cells (expressing a TCR specific for the LCMV glycoprotein gp33) were transduced with retroviral vector encoding hairpin RNAs (shRNAs) targeting our gene of interest (*Elf1* or *Prdm9*) or a negative control (*Cd19*), mixed at a 1:1 ratio then transferred into recipients that were subsequently infected with LCMV-Armstrong (Fig. 6a and Fig. S4a). On day 7 of infection, the effector P14 T cell populations in the spleen that were deficient for *Elf1* or *Prdm9* displayed an increased frequency of memory precursor (KLRG1^-^CD127^+^) cells compared to the control population (Fig. 6b,c). Furthermore, at day 21 of infection, the memory T cell population was comprised of an increased proportion of T_EM_ (CD62L^-^ CD127^+^) when *Elf1* expression was knocked down and an increased frequency T_CM_ (CD62L^+^CD127^+^) and a decreased frequency of t-T_EM_ (CD62L^-^CD127^-^) when *Prdm9* was knocked down (Fig. 6b,c). These data validate the functions of *Elf1* and *Prdm9* in CD8^+^ T cell differentiation and suggest a role for these TFs in promoting terminal differentiation.

**Fig. 6.**
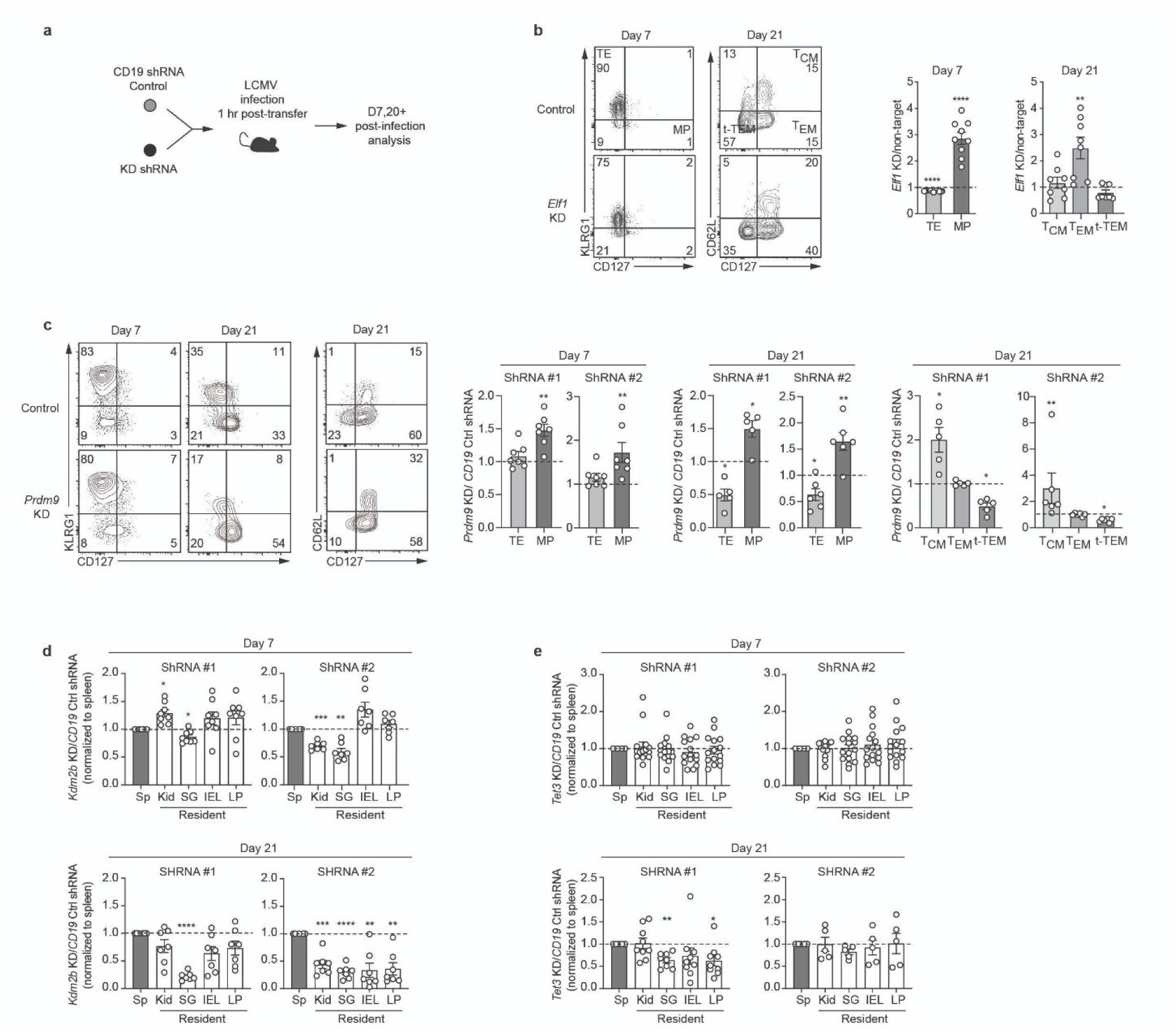
Validation of novel TFs’ roles in CD8^+^ T cell differentiation. **(a)** Experimental schematic in which congenically distinct P14 were transduced with gene-specific (knock-down, KD) or *Cd19* control shRNA encoding retroviruses, mixed 1:1 then transferred into recipient mice subsequently infected with LCMV. **(b)** The ratio of TE (KLRG1^hi^CD127^lo^) or MP (KLRG1^lo^CD127^hi^) and T_CM_ (CD62L^hi^CD127^hi^), T_EM_ (CD62L^lo^CD127^hi^) or t-T_EM_ (CD62L^lo^CD127^lo^) (gating strategy, left) within the *Elf1* KD and control P14 population recovered from the spleen on day 7 (middle) or day 21-27 (right) of infection. **(c)** Representative flow cytometry staining (left) and quantification of the ratio of TE and MP and T_CM_, T_EM_ or t-T_EM_ (right) within the *Prdm9* KD and control P14 population recovered from the spleen on day 7 or day 21. **(d)** The ratio of recovered *Kdm2b* KD and control P14 cells in indicated tissues on day 7 (top) and 21 (bottom) of infection. **(e)** The ratio of recovered *Tet3* KD and control P14 cells in indicated tissues on day 7 (top) and 21 (bottom) of infection. Data are cumulative of 2-3 experiments with n=3-5 per group. Graphs show mean ± SEM. A one-sample two-tailed t-test, two-tailed ratio paired t-test (B,C) or a one-way paired ANOVA test (D,E) was used to determine statistical significance; *p< 0.05, **p < 0.01, ***p < 0.001, ****< p < 0.0001.

Tissue-resident lymphocytes are a diverse population of non-recirculating lymphocytes lodged in tissues that provide localized protective immunity and immunosurveillance[74]. As the importance of this population to host protection is clear, we next sought to define regulators important for tissue residence. To identify putative tissue-residency-specific TFs, we focused on 6 lymphocyte populations originating from the small intestine (ILC3.NKp46+.SI, ILC3.NKp46-CCR6-.SI, ILC3.CCR+.SI, ILC2.SI, Treg.4.FP3+.Nrplo.Co, T8.IEL.LCMV.d7.SI) with 8 similar cells in the circulation as background (T8.Tcm.LCMV.d180.Sp, T8.Tem.LCMV.d180.Sp, T8.MP.LCMV.d7.Sp, T8.TE.LCMV.d7.Sp, Treg.4.25hi.Sp, NK.27+11b-.Sp, NK.27+11b+.Sp, NK.27-11b+.Sp). More than 60% of the identified TFs have been identified as regulators within the tissue-resident lymphocyte populations by previous studies (Table S3). For instance, *Hic1* is critical for establishing and maintaining T_RM_ cells in the gut[75,76], and *Ahr* is required for long term persistence of T_RM_ in the epidermis[77].

Using this approach, we also identified several genes with undefined function in tissue residency. We validated the role of 2 of these genes, *Kdm2b* and *Tet3*, in promoting differentiation of CD8^+^ T_RM_. Kdm2b and Tet3 both were both documented in the CIS-BP database to recognize specific DNA motifs and were thus included in our study. As above, congenically distinct P14 T cells expressing shRNAs for our gene of interest (*Kdm2b* or *Tet3*) or for a negative control (*Cd19*) were mixed 1:1 and transferred into recipients that were infected with LCMV-Armstrong (Fig. 6a and Fig. S4a). Circulating T cell populations of the spleen were compared to T_RM_ the salivary gland (SG), kidney (Kid), and small intestine intraepithelial layer (IEL) or lamina propria (LP) on day 7 and 21 of infection. By day 21, T cells lacking *Kdm2b* resided at a reduced frequency in all tissues examined compared to control cells (Fig. 6d). Furthermore, the proportion of T cells in the gut expressing the canonical T_RM_ markers CD69 and CD103 was reduced with decreased *Kdm2b* expression compared to control cells (Fig. S4b). *Tet3*-deficient T cells were also impaired in their ability to populate the SG and LP (Fig. 6e) at day 21 and showed an early defect in differentiation with reduced CD69 and CD103 expression on day 7 of infection compared to control IEL and LP T cells. Thus, the TFs, *Kdm2b* and *Tet3*, function to promote generation of CD8^+^ T_RM_ populations and represent newly identified regulators of T_RM_ identified regulators of this important T cell memory population, further supporting the tremendous predictive value of Taiji.

### Systematic comparison between human and mouse immune cells reveal conserved regulatory code

To systematically compare the regulatory code, we used Taiji to construct regulatory networks in human immune cell populations [16,17] similar to those we generated in the mouse immune system and obtained a PageRank matrix. Twenty-four human samples were paired with 29 mouse samples. These 53 samples spanning 8 immune cell lineages were used for all the downstream analysis in Fig. 7.

**Fig. 7.**
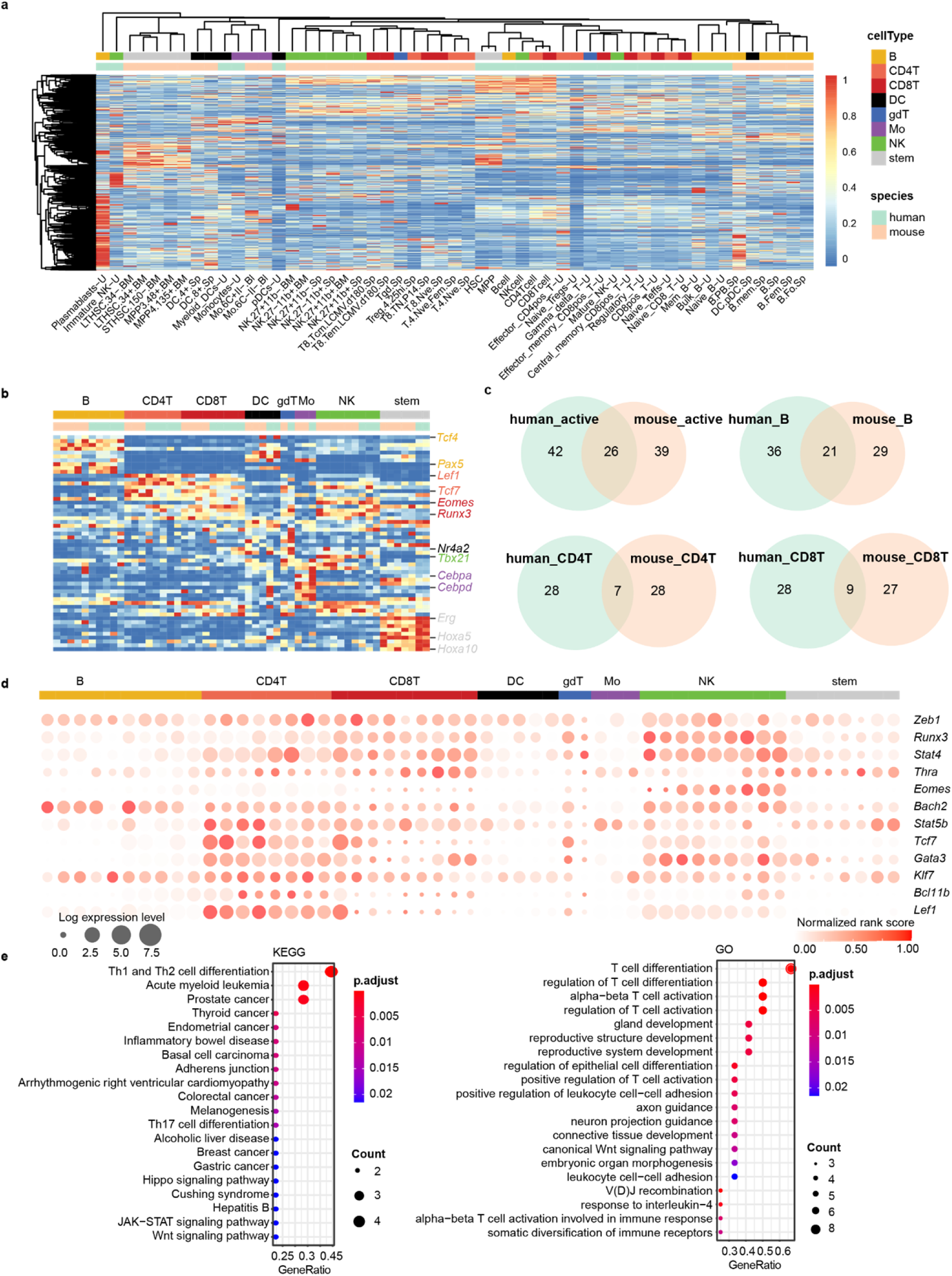
Comparison between human and mouse immune system. **(a)** Hierarchical clustering of 53 cell types from both human and mouse systems based on row-wise normalized PageRank scores of 733 TFs. TFs in rows, cell types in columns, and color of the cell in the matrix indicate the normalized PageRank scores with red displaying high scores. **(b)** PageRank scores heatmap of top10 cell lineage-specific TFs. Selected gene names are colored based on their cell lineage specificity. Legends and columns are the same as in **(a)**. **(c)** Comparison of active TFs and 3 comparisons of specific TFs between human and mouse systems. “Active” means the constitutively active TFs, whose average PageRank scores across all the cell types were ranked top 10% with CVs<0.5. **(d)** Twelve TFs important in several cell lineages in both mouse and human. Circle size represents the logarithm of gene expression and the color represents the normalized PageRank score. Column order is the same as **(b)**. **(e)** Enriched KEGG pathways and GO terms of 12 putative driver TFs identified in **(d)**.

The hierarchical clustering based on PageRank scores of 733 common TFs showed that samples with similar cell identity are more likely to be clustered together (Fig. 7a). For instance, B cells from both mouse and human are clustered, as well as DCs and monocytes. CD4^+^ and CD8^+^ T cells show more complex patterns indicative of the cell lineage heterogeneity both within and across the source organisms. The identified cell-lineage specific TFs for human and mouse following the same method described previously in “Materials and Methods: Identification of putative driver TFs in cell lineage specification”. The comparison of specific TFs between human and mouse revealed similarTF regulatory profiles across species (Fig. 7c). Twenty-four percent of total B cell-specific TFs are shared between the two organisms including Tcf4[38–41], Pax5[78–80], Bhlhe41[81–83], and several Irf family TFs (Irf4, Irf5, Irf8)[24,84,85]. This relatively large overlapping percentage is consistent with the clustering of B cells across species shown in Fig. 7a. Similarly, the overlapping percent of constitutively active TFs is relatively high (24%), showing that these house-keeping genes are required for fundamental biological functions across a variety of cell types and species. On the other hand, the overlap percentages for CD4^+^ and CD8^+^ T cells are relatively small, 11% and 14%, respectively. The conserved TFs like Eomes[86–88], Bcl11b[89–91], Lef1 and Tcf7[27,92,93] are well-known for their important roles in T cell differentiation in both systems. We also identified species-specific TFs including Msc, Maz, Klf5 and Snai3, Sp4, and Foxo4 in human and mouse CD8^+^ T cells, respectively.

The heatmap in Fig. 7b displays the PageRanks of cell-lineage specific TFs shared between mouse and human. Some of them are well-known in both organisms, like Cebpd, Cebpa as monocytes-specific TFs[94,95]; Erg[96,97], several Hoxa[98–100] family TFs as stem cell-specific TFs. Some of these specific TFs can be very important in several cell lineages in both mouse and human systems, like Tcf7, Lef1, Bcl11b, Gata3 show in Fig. 7d, suggesting the potential sophisticated regulatory roles in closely-related cell lineages like CD4^+^ and CD8^+^ T cells. The functional analysis of TFs shows strong connection with regulation of T cell differentiation (Fig. 7e). This comparative analysis between human and mouse reveals some common regulatory code across species, which would be very valuable for elucidating the functions of unknown TFs in the human immune system.

## Discussion

The mouse immune system provides an ideal setting for implementing system-level analyses of cellular differentiation as well-defined cell states have been established and the differentiation stages from progenitor to mature cells have been thoroughly characterized[101]. Considerable efforts have been made in mapping the regulatory networks driving immune cell differentiation, yet identification of the critical TFs that regulate cell fate decisions at key differentiation branch points has remained a challenge. Here, we have systematically identified key TFs in 9 immune cell lineages that encompass a range of key lineages in the mouse and human immune systems based on the Taiji framework. Taiji integrates gene expression and epigenomic information (ATAC-seq or histone modification) to construct regulatory networks, thus better delineating the cell identity than single-omics data. Furthermore, Taiji calculates PageRank scores for each regulator by considering the impacts of its upstream regulators and downstream regulatees including the feedback from its regulatees. Importantly, Taiji considers not only the expression levels of a gene’s regulators and regulatees but also the strength of each regulatory interaction indicated by the extent of open chromatin, motif match, and expression of the TF itself. Therefore, the PageRank score of a gene in Taiji represents its predicted global importance in the genetic network. The “master regulators” would have the highest PageRank scores as they regulate many TFs that in turn regulate highly expressed genes. Note that the majority of the TFs have PageRanks scores well correlated with their expression levels but there are a substantial portion that show weak or no correlation. These TFs have moderate expression themselves but regulate many highly expressed genes and their regulatory activities are thus better characterized by PageRanks scores, which is a particularly powerful benefit of this approach: Taiji assesses functional regulation of target genes rather than TF expression dynamics.

Other similar methods that identify important TFs have been reported before. Specifically, SCENIC developed by Aibar et al 2017[8] aims to analyze single cell RNA-seq data without considering ATAC-seq. This method only uses gene expression data and constructs the genetic network by predicting each gene’s expression using the TFs’ expression levels and finding the most predictive TFs as the regulators. SCENIC cannot be applied to the bulk mouse immune data[15] as Taiji did and does not have enough samples to select the regulator TFs. Furthermore, the motif analysis in SCENIC only considers sequences around TSS while Taiji includes both promoters and enhancers.

Importantly, Taiji integrates RNA-seq and ATAC-seq data to construct a genetic network so it can capture the global impact of TFs in the genetic network. Taiji considers not only the effect on TF’s direct target genes, but also all the descendants in the whole network. Due to the iterative calculation of PageRank scores, Taiji takes into account the impact of its upstream regulators and downstream regulatees including the feedback from its regulatees, which makes Taiji capable of assessing the global influence for individual TFs. This global view is what distinguishes Taiji from other methods. It is worth noting that we have previously shown that Taiji is very robust and resistant to noise[12] (Fig. 2 in Zhang et al., 2019). The PageRank scores can correctly predict TF’s importance even when 80% of the edges in the network are disturbed.

As for the comparison with the Maslova et al. 2020[9] analysis, Taiji integrates RNA-seq and ATAC-seq to construct a genetic network and assess the global importance of TFs while AI-TAC focuses exclusively on predicting ATAC-seq signals using sequences and extracting the motifs predictive of cell type specification. Maslova et al. did not use gene expression information in their analysis and did not build a genetic network. Identifying enriched motifs in specific cell types is equivalent to considering the direct targets of TFs when evaluating the TF importance. While such an approach is useful, we have demonstrated in our previous work[12,13] that the PageRank scores performed better by considering not only the direct target genes of TFs but also their own regulators and the descendant genes of their direct targets. In this sense, our study is the first systems biology approach that integrates RNA-seq and ATAC-seq to systematically identify the key regulators during mouse immune system development.

Furthermore, if a motif is weak, the PageScore of the TF is decided by whether these motifs occur in the open chromatin regions (measured by the peak intensity of the ATAC-seq data), the TF expression and its target expression levels. Because we also compare TFs between different cell types to identify cell-type/lineage specific regulators. The relative difference between the PageRank scores of TFs also helps to uncover important TFs with weak motifs. As such, our method identifies GATA3 as an important TF in several T cell subsets (see Table. S2), which is a CD4^+^ T cell master regulator with weakly defined sites. Therefore, integration of motif strength with open chromatin strength, expressions of TF and its target genes is superior to only considering motifs that are present or absent in open chromatin as in Maslova et al. study.

Another advantage of Taiji is its capability to construct genetic networks in individual cell types and thus allows identifying cell-type specific regulators. By comparing PageRank scores of TFs across cell types, we have successfully identified cell lineage-specific putative driver TFs, thus constructing a comprehensive catalog of key regulators in various immune cell types. In addition to identifying dozens of known regulators, for instance, *Tcf7*[32–36], *Lef1*[27,33,34], *Stat4*[102], *Gata3*[103] as T cell-specific TFs; *Bcl11a*[21,22], *Tcf4*[38–41], *Ebf1*[79,104,105], *Pax5*[78–80] etc. as B cell-specific TFs, we have also found several TFs with previously undefined roles in regulation of immune cell lineage differentiation and development, for instance, *Pbx4* may have important role in T cell development; *Meis1*, *Jdp2* may be critical regulators in ILC lineage development. Overall, the identified novel cell type-specific TFs provide insights for the future experimental validation to fully reveal the underlying regulatory mechanisms.

We also provided here a comprehensive temporal transcriptomic and epigenomic dynamics in a diverse set of cell types, revealing “transcriptional waves” in which combinations of TFs are particularly active in specific cell types during specific developmental stages. The observed transcriptional waves patterns suggest the possible mechanisms of how the TF activities are coordinated to orchestrate the cell type differentiation. We found different types of transcriptional waves: constitutively active waves (Fig. 5a,b), cell lineage-specific waves (Fig. 5c,d), and stage-specific waves (Fig. 5e-h). The cell lineage-specific waves are composed of coordinately highly ranked TFs across multiple cell types in the same cell lineage. This indicates that certain TFs are “multi-taskers”[106] which play critical roles in multiple or successive developmental stages along one lineage. The stage-specific waves, on the other hand, show TF combinations having exclusively important roles during a very specific developmental stage. For example, three *Duxbl* family members display high rank in preT.DN3.Th (Fig. 5f), consistent with that *Duxbl* was shown to have high recombination activity before β-selection and result in a developmental block at the DN3-to-DN4 transition due to enhanced apoptosis and reduced proliferation of pre-T cells in thymus[60]; noticeably, there are additional TFs in the same cluster, such as *Lcor*, *Tcf4* and *Runx2*, suggesting their possible roles in the development of preT.DN3.Th. Another example is C10, which is specifically important in T8.Tem.LCMV.d180.Sp and most of the 19 TFs have not been reported to function in T_EM_, such as *Rfx6*, *Lhx4* and *Atf6*. Identification of these transcriptional waves with defined regulatory patterns and predicted regulatory interactions not only uncover new regulators but also would greatly facilitate mechanistic studies.

To reveal the conserved regulatory code across organisms, we also constructed the regulatory networks in matched human immune cells using Taiji framework. The systematic comparison of constitutively active TFs and cell lineage-specific TFs showed that some TFs undertake similar crucial roles in the development and differentiation of immune cells in both human and mouse systems, like *Stat1*, *Stat3*[107–109], *Sp2* and *Sp3*[110–112] as common constitutively active TFs; *Eomes*[86–88], *Bcl11b*[89–91], *Lef1* and *Tcf7*[27,92,93] as CD4^+^ and CD8^+^ T-important TFs. This systematic analysis that spans all the immune cell lineages identifies the conserved TFs across species and shows the potential of elucidating the molecular functions of unknown TFs in human immune system.

To demonstrate the power of our analysis, we experimentally validated 4 regulators that have no reported functions in circulating and tissue-resident CD8^+^ T cell memory. T cells responding to infection engage a transcriptional network that directs differentiation of progeny that adopt a range of cellular states; thus, creating a heterogeneous population of effector T cells composed of cells able to eliminate the present infection and others capable of providing robust long-lasting immunological memory[106,113]. A large fraction of effector T cells are terminally differentiated and after providing immediate effector function to clear the pathogen from the host will undergo apoptosis. Alternatively, other cells will be maintained in a ‘stem-like’ state capable of seeding the memory T cell population that provides protection from reinfection[106,113]. The T cells within the memory population also exist in a spectrum of phenotypes. T_EM_ and T_RM_ reside in the circulation and peripheral tissues, respectively, and provide an immediate effector response while T_CM_ remains less differentiated, able to undergo extensive proliferation to generate a new burst of effector T cells[106]. While key transcriptional regulators that program these cellular fates have been identified[114], a comprehensive network of TFs has not been completed, particularly when the subsets represented within the memory T cell population are considered. Our analysis uniquely compared several memory B and T cell populations as a whole to effector populations and identified a number of novel putative important TFs of memory states, including *Elf1* and *Prdm9*. Elf1 has been associated with NKT cell development[115] and Prdm9 with regulation of cell cycle, a process closely linked with cellular differentiation[116,117]. Our validation of their role in CD8^+^ T cell differentiation following acute infection clearly demonstrated essential roles for these TFs in promoting the terminally differentiated state of CD8^+^ effector T cells. This suggests that *Elf1* and *Prdm9* control the terminal differentiation transcriptional program directly or inhibit the program that promotes formation of the memory cell state. While our analysis predicts relative regulatory weight of the TF at the given cell-fate branch point, it cannot determine direction of regulation (ie. whether it promotes or inhibits). Thus, additional hits from our analysis may be vital for regulating differentiation of the memory T cell subsets as suggested by the identification of the well-known TF *Foxo1* that promotes and sustains the CD8^+^ T cell memory population[66–69].

While distinct regulatory networks drive the differentiation and maintenance of effector and memory CD8^+^ T cells cell states, additional transcriptional programs define the trafficking patterns and location of T cells within the host. To identify TFs important for directing the residency of T cells within peripheral tissues, we compared several lymphocyte populations localized to the gut to those found in the circulation. We identified numerous known and also unknown regulators including Kdm2b and Tet3. We validated the function of *Kdm2b* and *Tet3* in the formation of CD8^+^ T_RM_ across several tissues. CD8^+^ T_RM_ failed to accumulate the following infection in the absence of *Kdm2b*. KDM2B has been shown to promote not only cell proliferation but also migration of mouse embryonic fibroblasts[118] and analogous mechanisms may be in play in tissue-resident lymphocyte populations. Our validation experiments also revealed that the differentiation of CD103^+^ T_RM_ cells were delayed with the loss of Tet3. Interestingly, Tet3 has been linked to the regulation of multiple key TGF pathway genes[119], a signaling pathway that is critical for the generation and maintenance of CD103^+^CD8^+^ T_RM_[120].

Taken together, our analyses have identified critical TFs that regulate two important T cell differentiation pathways - determination of cell state/fate and cellular trafficking/localization. The literature evidence of well-known TFs and the validation experimental results of novel regulators suggest the power of our study in guiding mechanistic investigations in the future. With the increasing understanding of CD8^+^ T cell community, our approach will also be applicable to further identify key regulators in the finer definition of CD8^+^ T cells, thus facilitating the novel discovery of CD8^+^T cell diversities and functions. Importantly, our approach is generally applicable to any cell type. Given the fast accumulating transcriptomic and epigenomic data generated from projects such as Human Cell Atlas, Taiji provides a powerful tool to profile the landscape of regulators that define cell-type and cell-state specificity, which would facilitate understanding the underlying mechanisms and elucidate manipulations such as overexpression of the key regulators to manipulate cell fate.

## Acknowledgements

This work was partially supported by the NIH (R01AI150282 and P01AI132122).

## Author contributions

C.L. designed studies, performed all the computational analysis, prepared the figures and tables, wrote the manuscript; K.O. designed and performed experiments, analysed the data, prepared the figure and wrote the manuscript; C.T. and N.K. performed experiments, analysed the data and prepared the figure; J.T.C. contributed advice and supervised the experiment; W.W. and A.W.G. supervised the project, designed studies, and wrote the manuscript.

## Supplementary Figures

**fig. S1.**
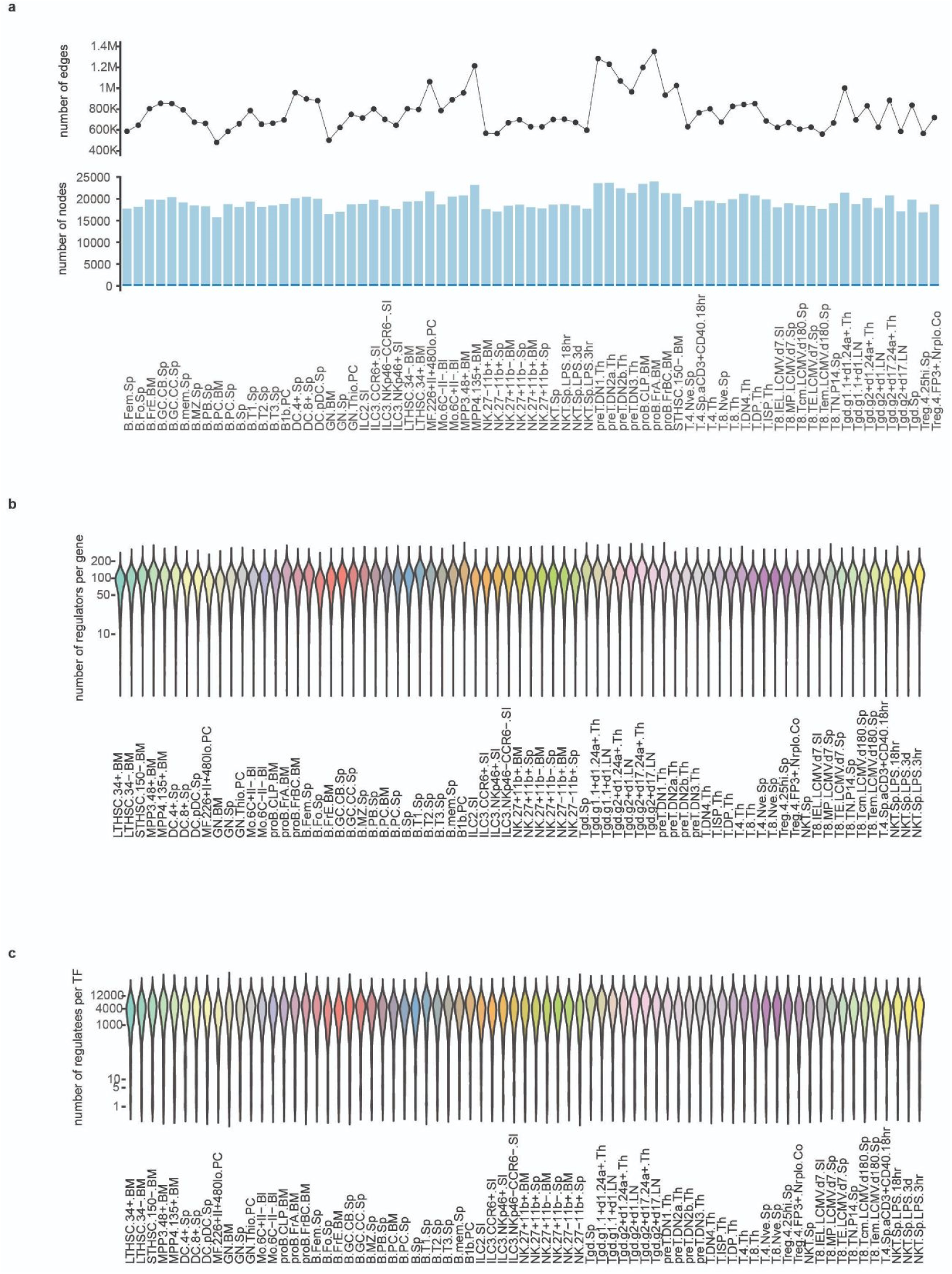
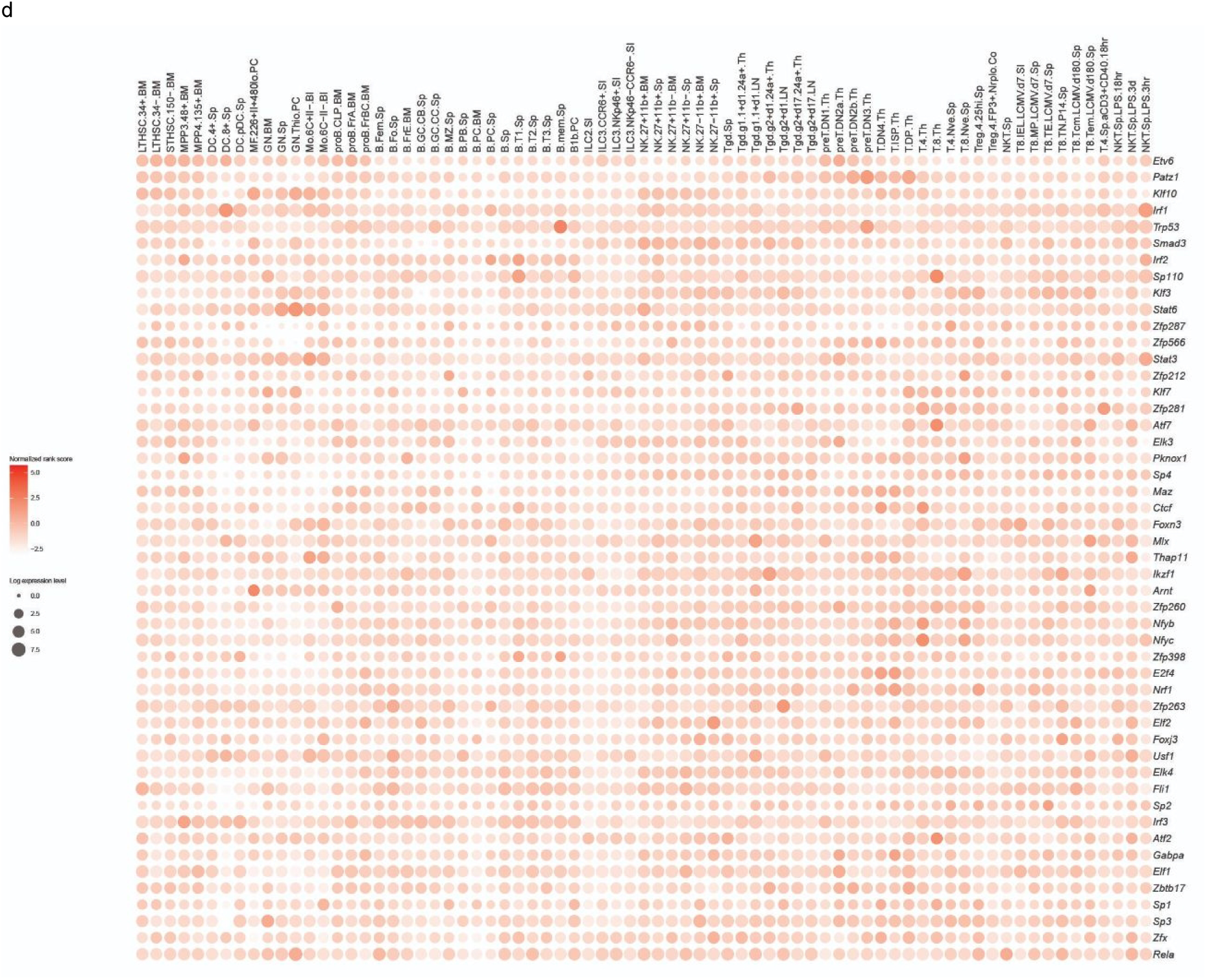
Topological properties of genetic networks. **(a)** The number of edges (top) and nodes (bottom) in each genetic network. **(b)** The distribution of the number of regulators per gene for each genetic network. **(c)** The distribution of the number of regulatees per TF for each genetic network. **(d)** PageRank scores (color shade) and expression levels (circle size) of 48 constitutively active TFs across all 73 cell types.

**fig. S2.**
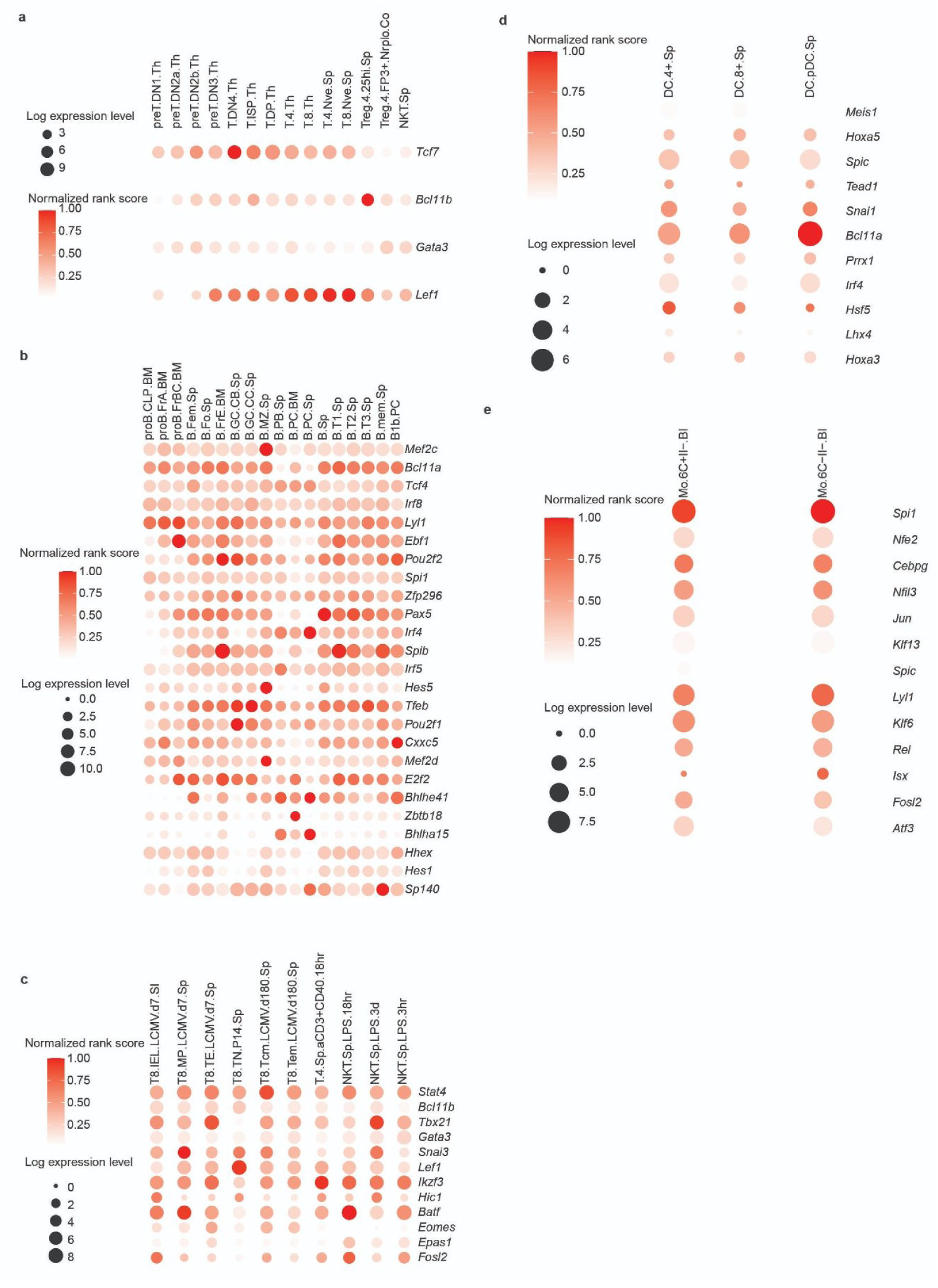
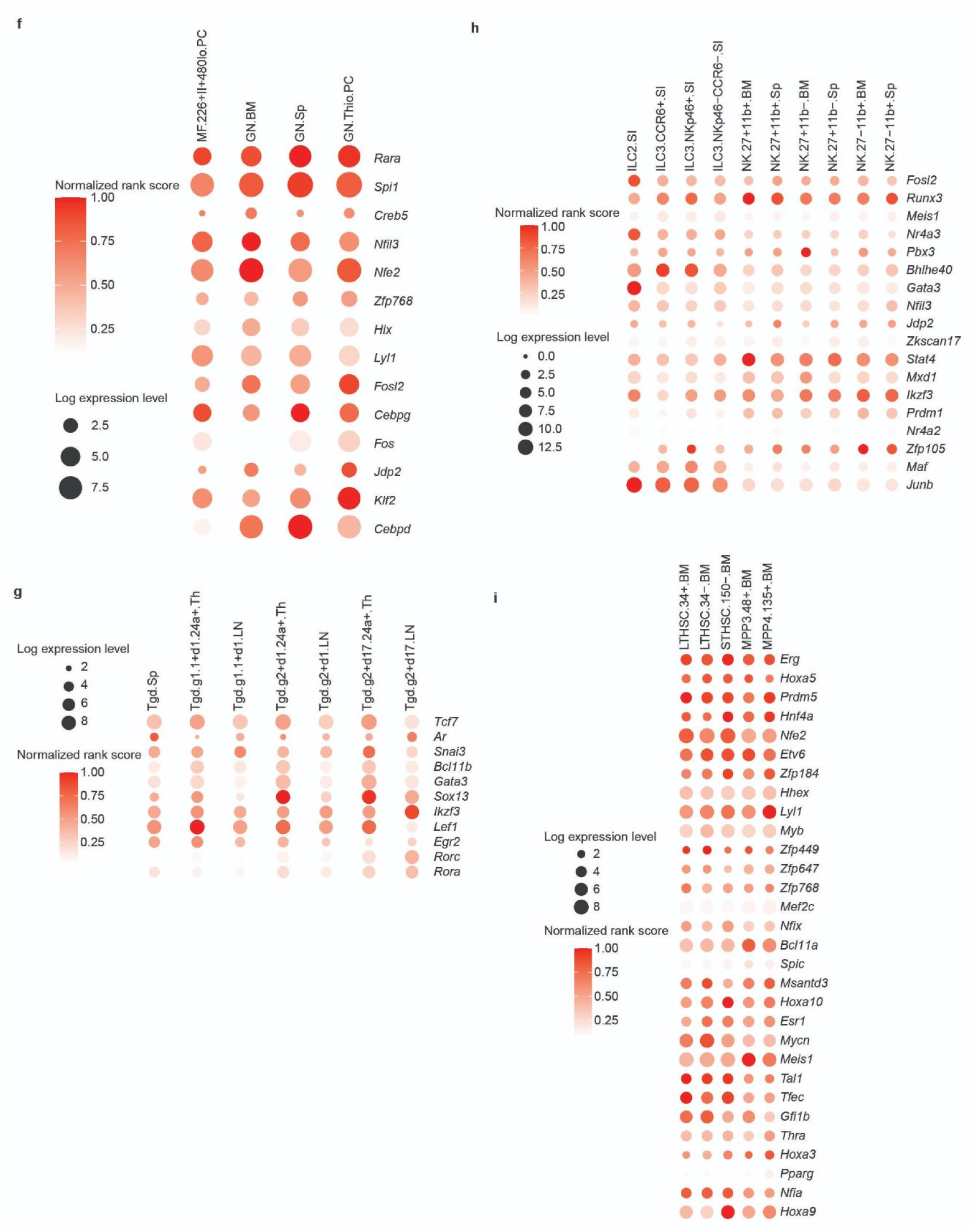
Identification of important TFs in 9 immune cell lineages. **(a)** αβT cell. **(b)** B cell. **(c)** Act_T cell. **(d)** DC cell. **(e)** Mo. **(f)** MF/GN. **(g)** γδT cell. **(h)** ILC. **(i)** Stem.

**fig. S3.**
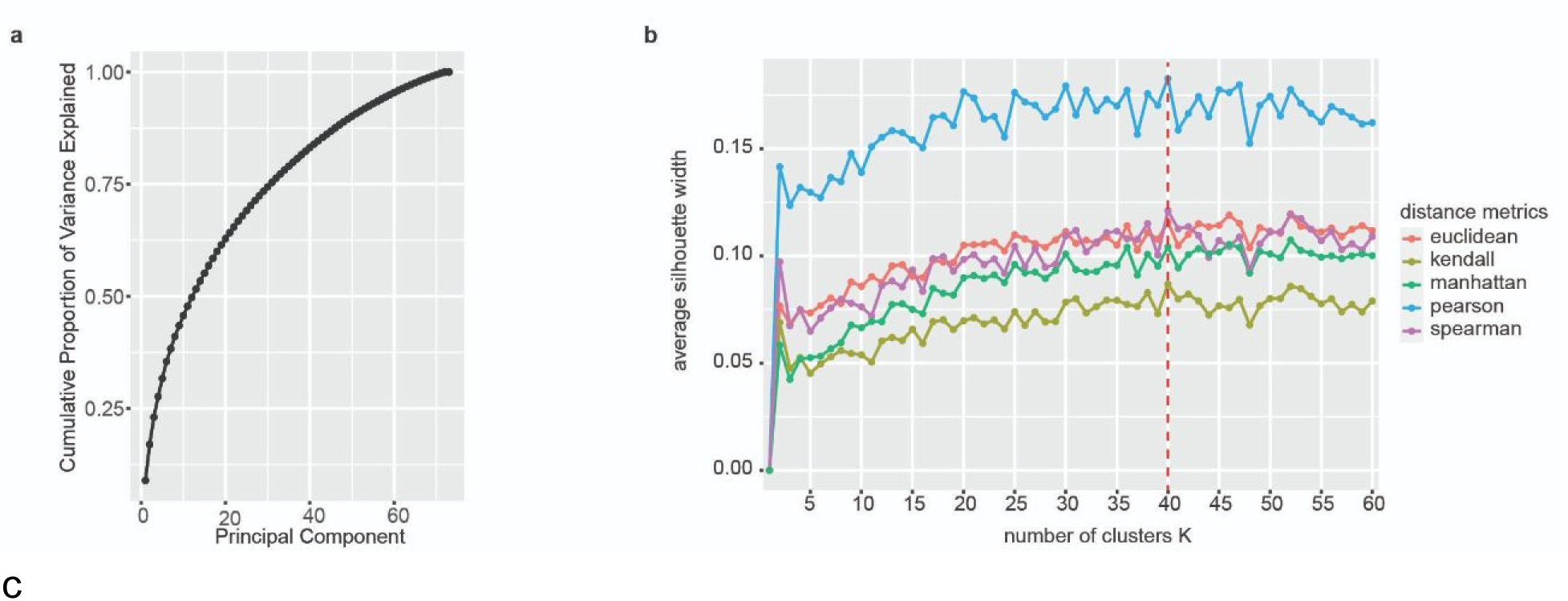
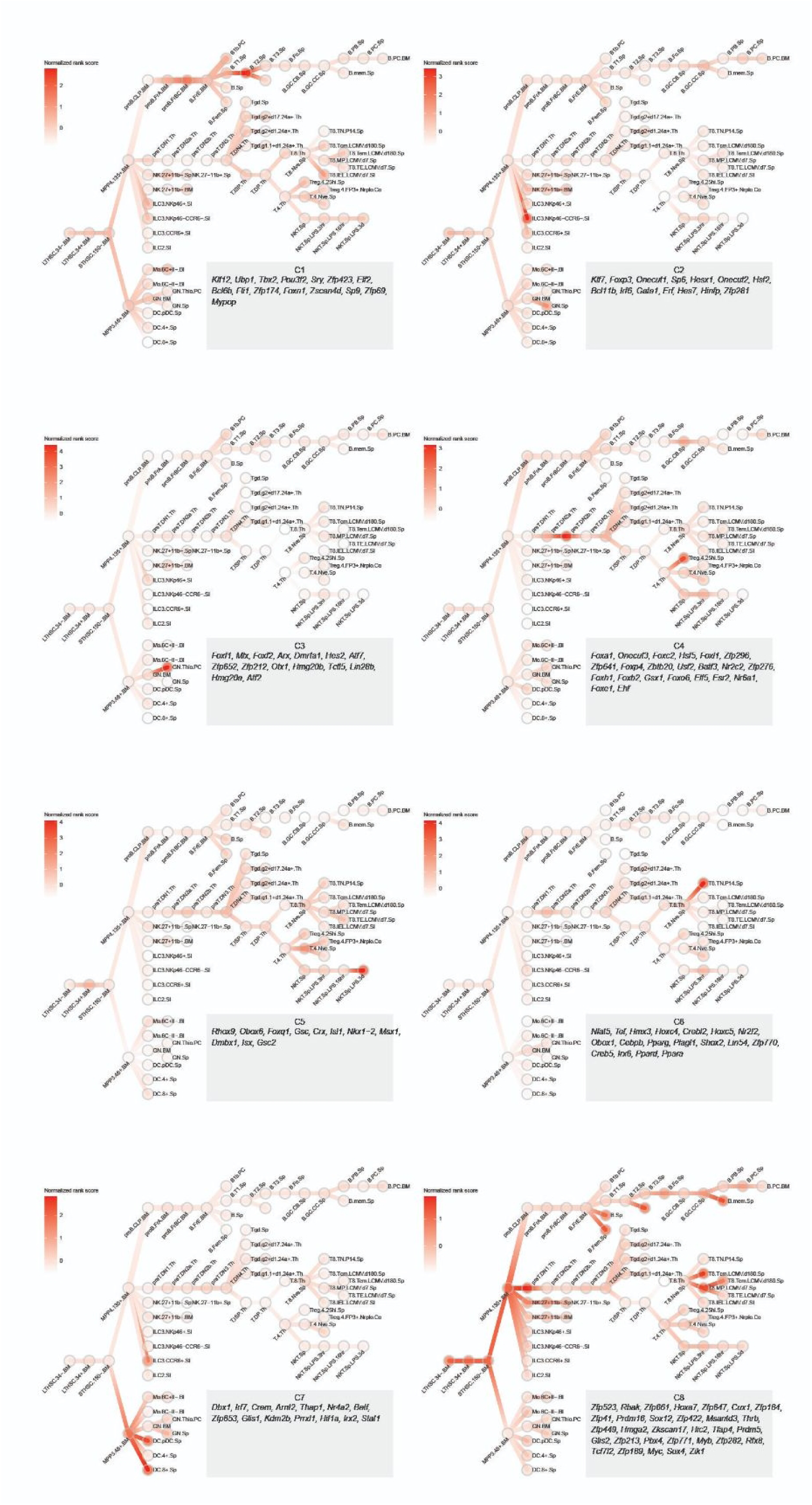
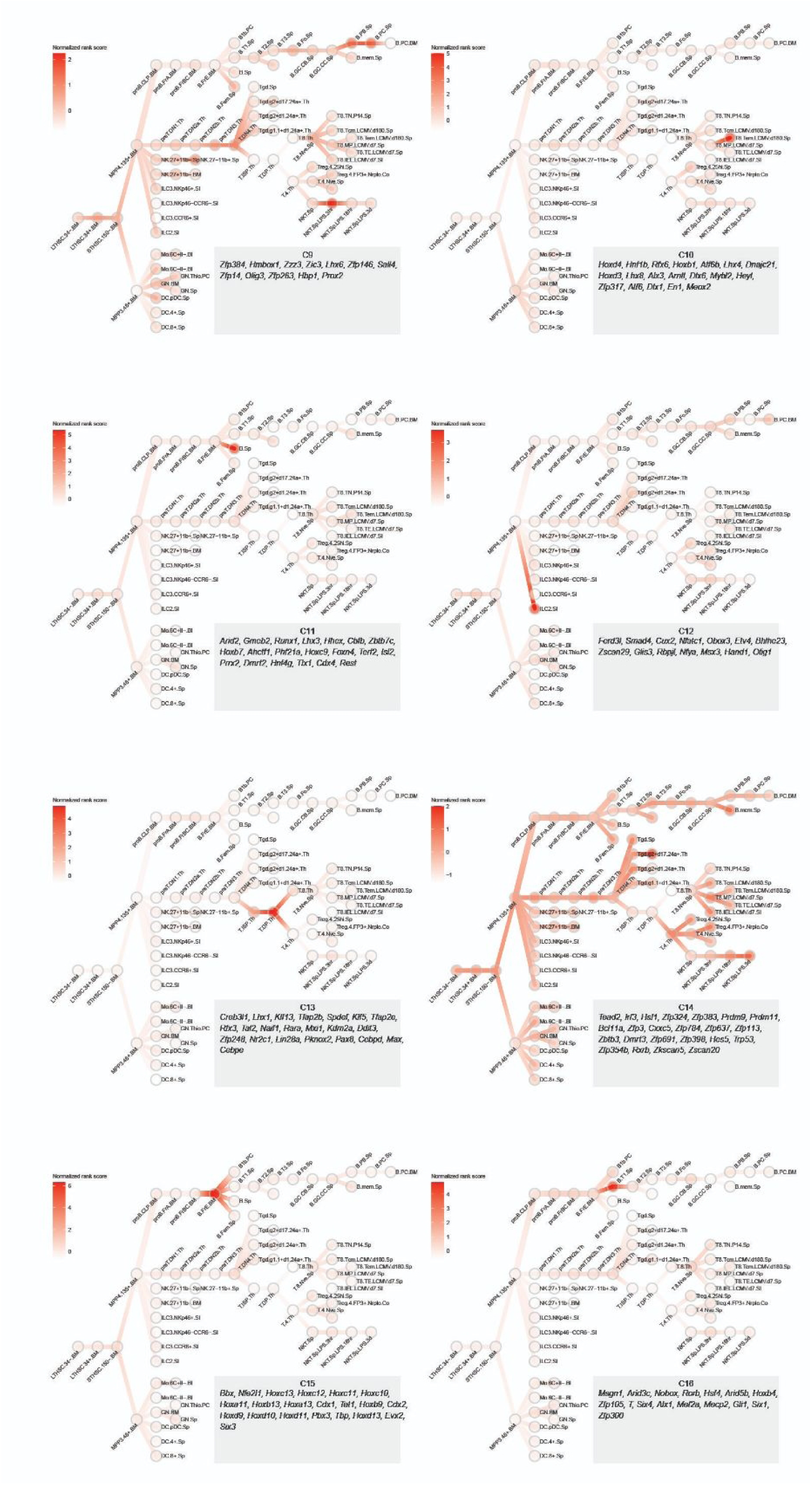
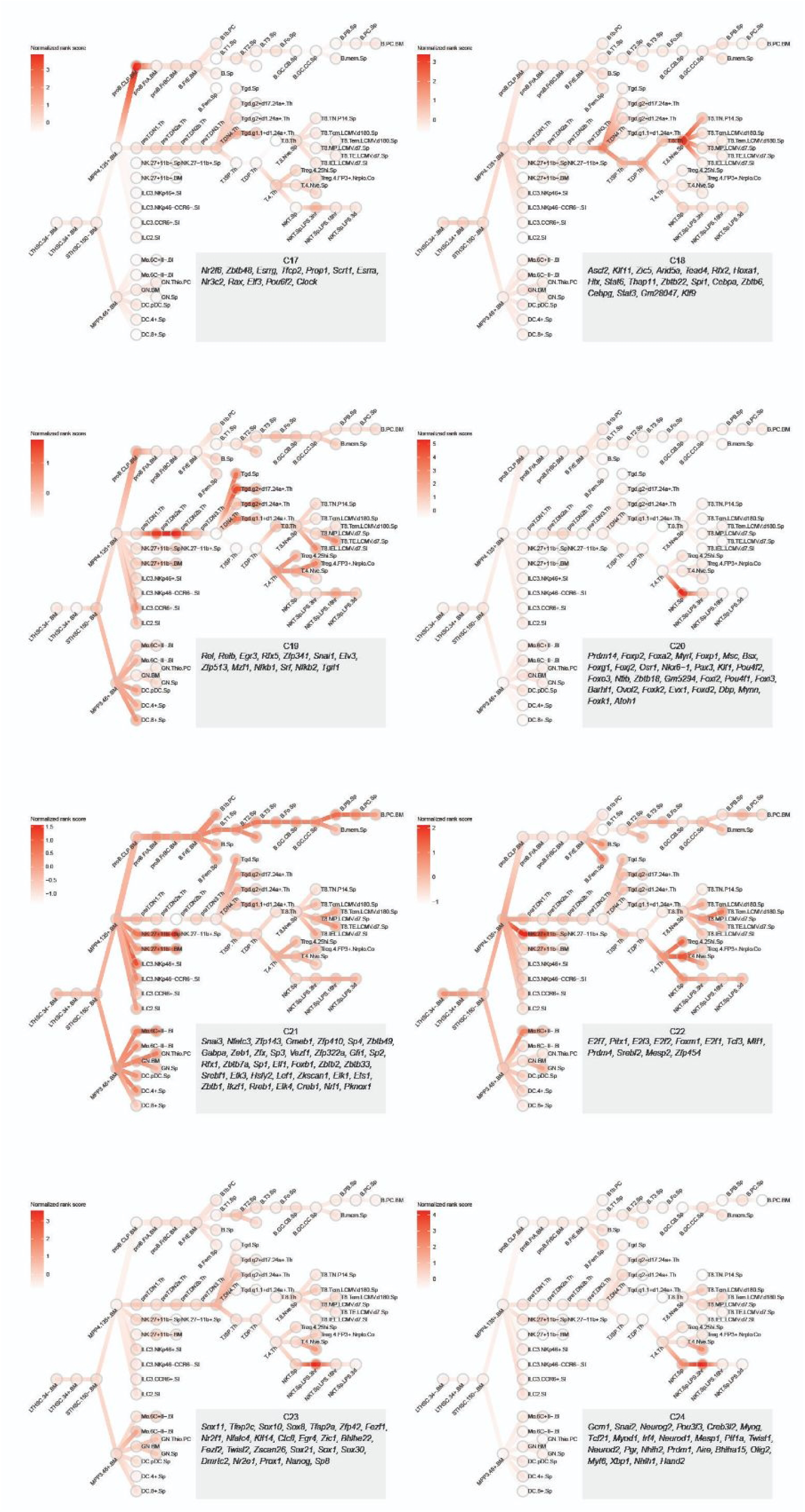
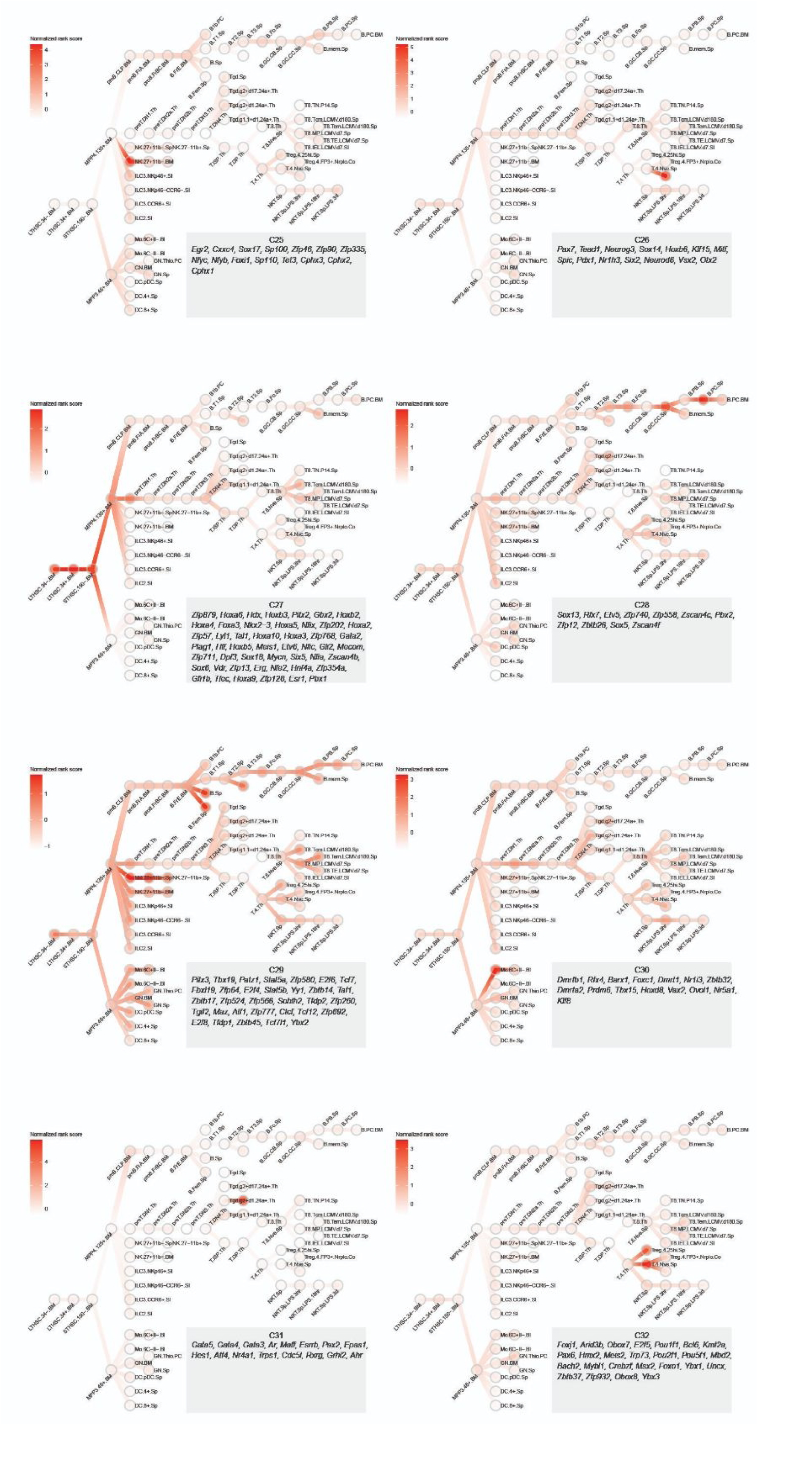
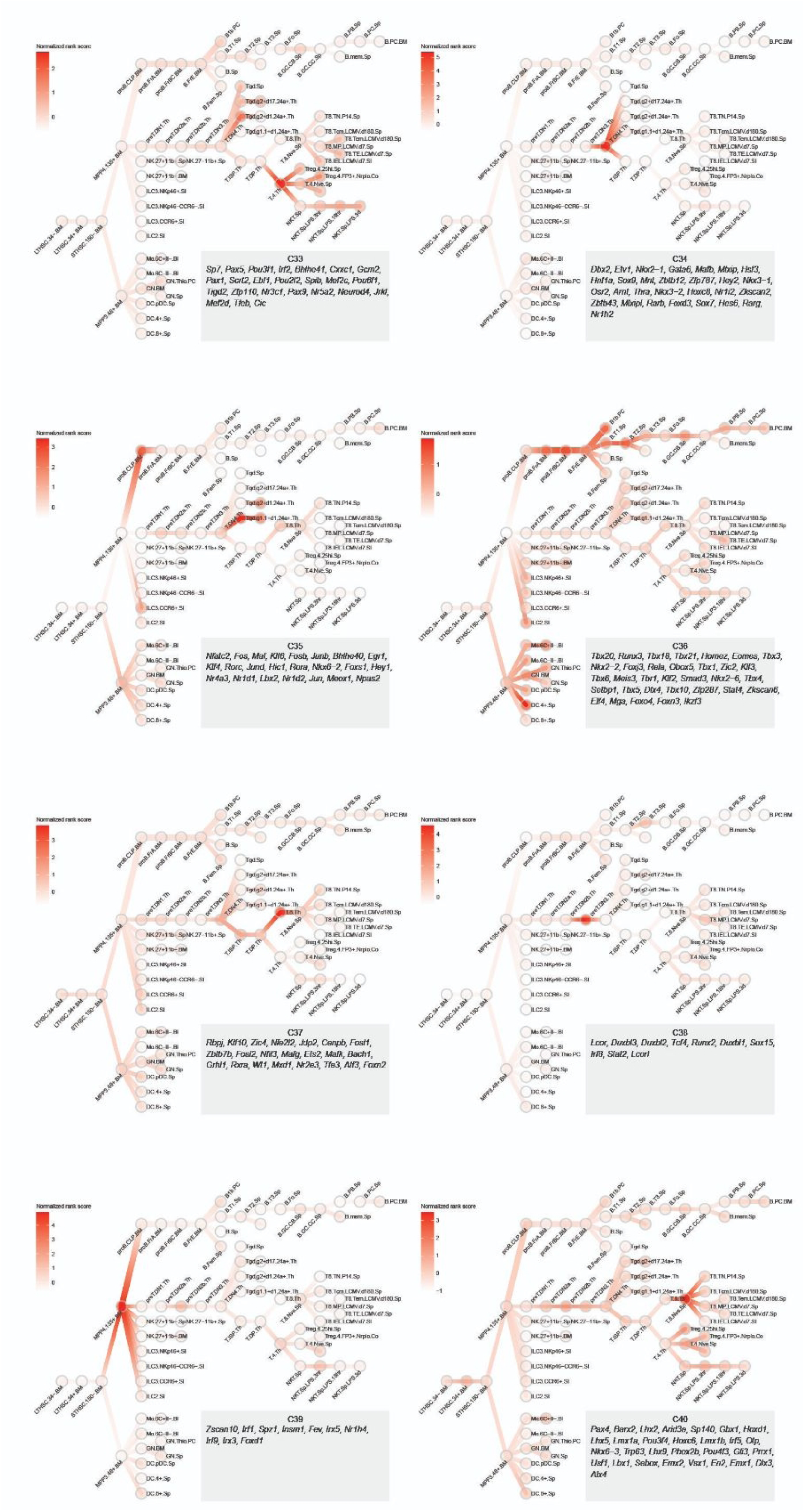
Transcriptional waves direct cell lineage differentiation in the mouse immune system. **(a)** Plotting the cumulative proportion of variance explained against the number of principal components (PCs). The first 30 PCs were kept according to the “elbow” method. **(b)** Selecting the best distance metric and number of clusters according to the Silhouette metric. The Pearson correlation was chosen and k=40 was the ideal cluster number. (**c**) 40 transcriptional waves with clusters of TFs.

**fig. S4.**
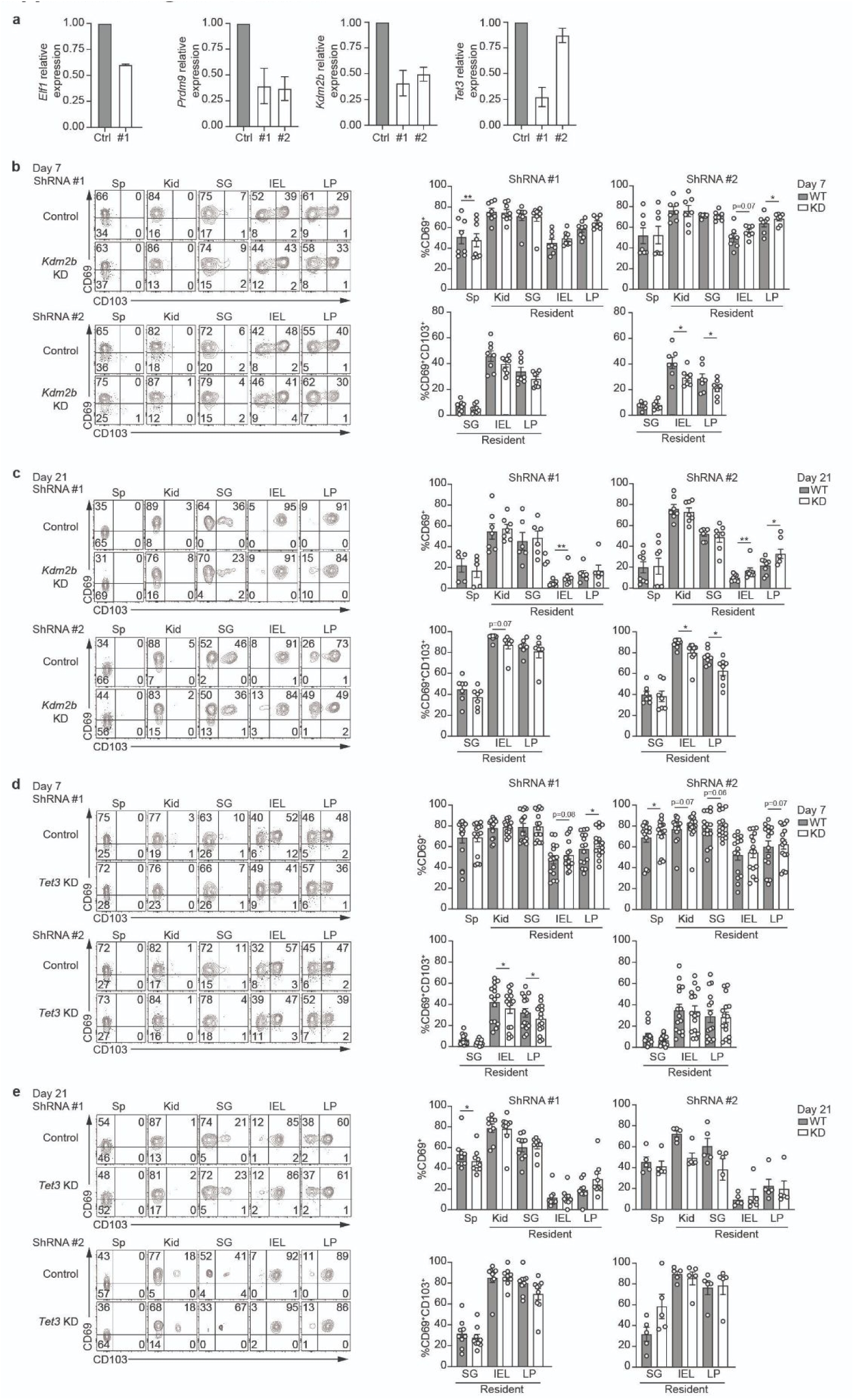
Validation of novel TFs’ roles in CD8^+^T memory cell formation and tissue residency. **(a)** qPCR analysis to assess knockdown efficiency of indicated gene in P14 cells prior to transfer. **(b,c)** Representative expression of CD69 and CD103 on *Kdm2b* KD and control P14 cells recovered from indicated tissue on day 7 and 21 of infection (left). Quantification of the frequency of populations (right). **(d,e)** Representative expression of CD69 and CD103 on *Tet3* KD and control P14 cells recovered from indicated tissue on day 7 and 21 of infection (left). Quantification of the frequency of populations (right). Numbers in graphs indicate percent of cells in the corresponding gate. Data are cumulative of 2 (b,c) or 3 (d,e) independent experiments with n=3-4. Graphs show mean ± SEM. A two-tailed ratio paired t-test was used to determine statistical significance; *p< 0.05, **p < 0.01.

## Supplementary Tables

**table S1.**
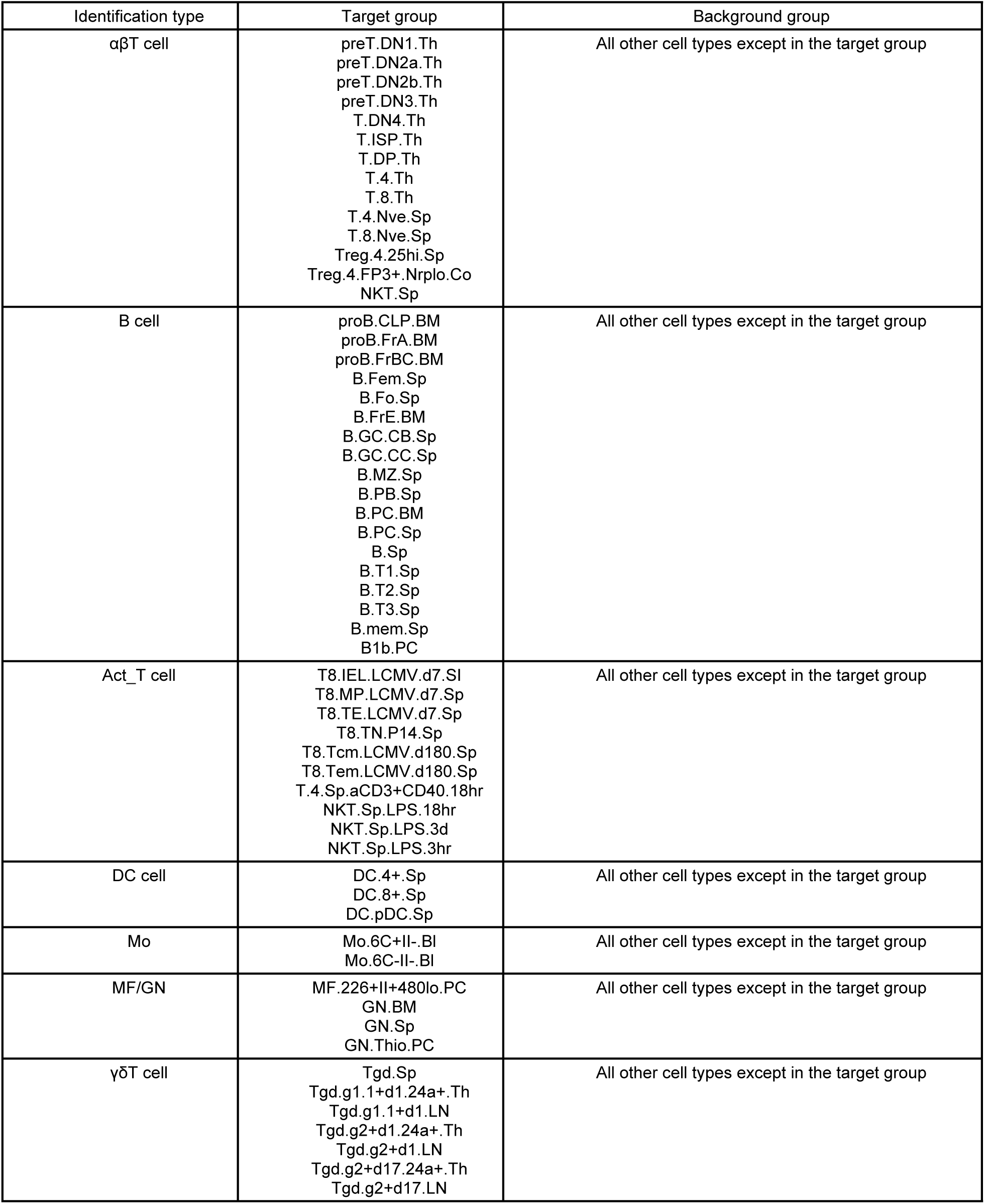
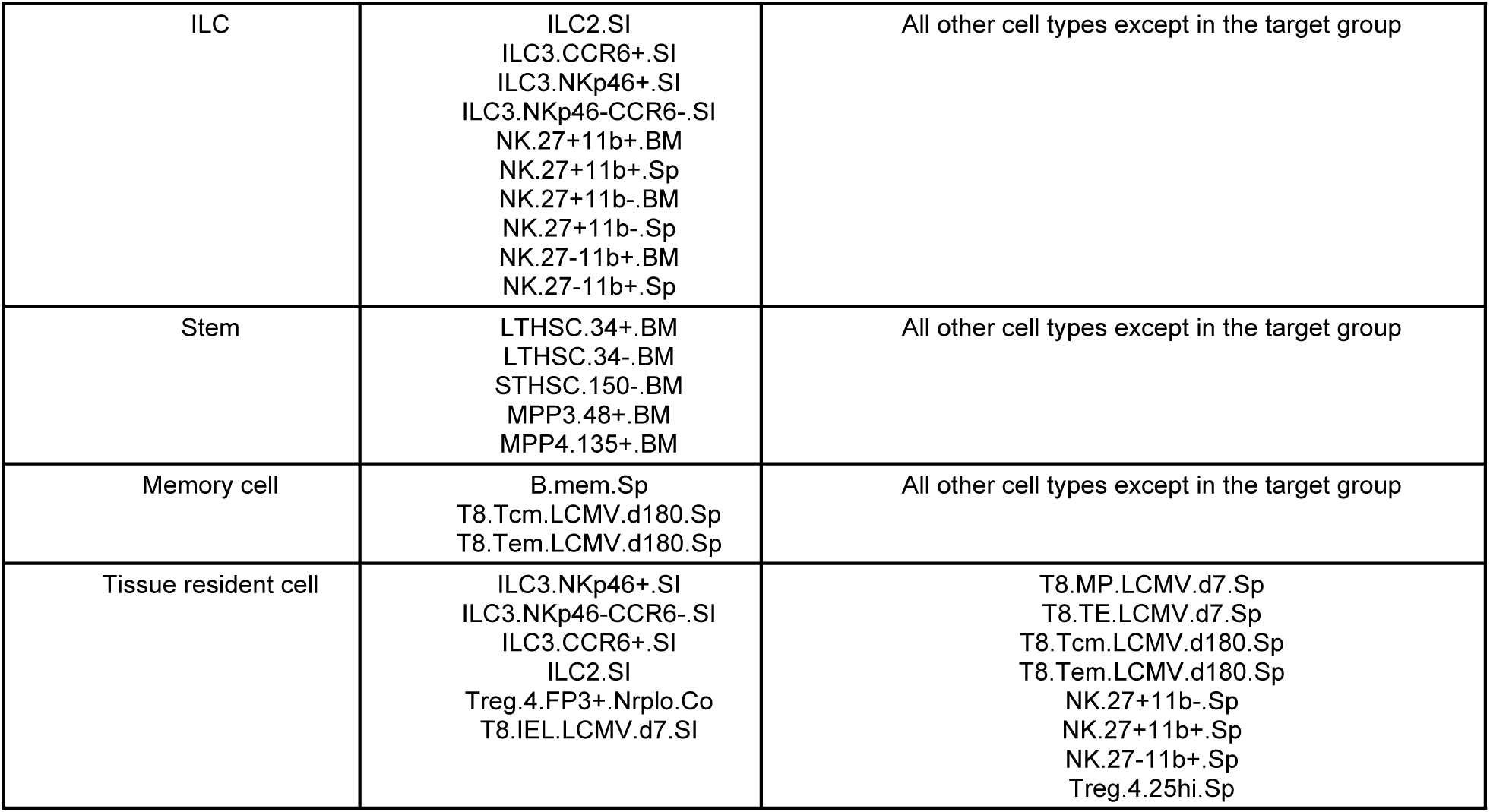
Cell types in target/background groups in the identification of putative driver TFs.

**table S2.** putative driver TFs in the five cell lineages.

B cell
| Predicted important TFs | Evidence |
| --- | --- |
| *Mef2c* | Essential for B cell early development[121] |
| *Bcl11a* | Relates to B cell development[22] |
| *Tcf4* | Essential for B cell early development[41] |
| *Irf8* | Regulates to B cell differentiation[24] |
| *Lyl1* | Induces B-cell lymphoma[122] |
| *Ebf1* | Essential for B cell early development[104] |
| *Pou2f2* | Regulates B cell differentiation[123] |
| *Spi1* | Essential for B cell differentiation[124] |
| *Zfp296* | Unknown |
| *Pax5* | Essential for B cell early development[78] |

αβT cell
| Predicted putative driver TFs | Evidence |
| --- | --- |
| *Tcf7* | Relates to αβT cell differentiation[125] |
| *Bcl11b* | Required for αβT cell differentiation and survival[126] |
| *Gata3* | Essential for CD4<sup>+</sup> T cell differentiation[103] |
| *Tgif2* | TGF-β–signaling components[126] |
| *Zbtb1* | Essential for development of T cells[127,128] |
| *Lef1* | Relates to αβT cell differentiation[129] |
| *Zfp566* | Unknown |
| *Stat5b* | Promotes the expansion of mature $\alpha\beta$T cells[130,131] |
| *Zfp64* | Promotes Toll-like receptor-triggered innate immune response[132,133] |
| *Stat5a* | Indispensable in Treg cell development[131,134] |

Act T cell
| Predicted putative driver TFs | Evidence |
| --- | --- |
| *Stat4* | Required in CD4<sup>+</sup> T helper cell differentiation[102] |
| *Bcl11b* | Controls CD8<sup>+</sup> T cell fate decisions[89] |
| *Tbx21* | Orchestrate Th1, Th2, and CD8<sup>+</sup> T cell differentiation[135,136] |
| *Gata3* | Controls Th2, CD8<sup>+</sup> T cell proliferation[137] |
| *Snai3* | Regulate CD8<sup>+</sup> T cell activation[138] |
| *Lef1* | Regulate CD8<sup>+</sup> T cell differentiation[34] |
| *Ikzf3* | Controls CD8<sup>+</sup> T cell responses[71] |
| *Hic1* | Critical in T cell function regulation[76] |
| *Batf* | Essential checkpoint in early effector CD8<sup>+</sup> T cell[139] |
| *Eomes* | Relates to T cell activation[88] |

$\gamma\delta$ T cell
| Predicted putative driver TFs | Evidence |
| --- | --- |
| *Tcf7* | $\gamma\delta$T cell-specific gene[140] |
| *Ar* | Unknown |
| *Snai3* | Unknown |
| *Bcl11b* | Relates to $\gamma\delta$T cell development[141] |
| *Gata3* | Correlates with $\gamma\delta$T cell functions[142] |
| *Sox13* | $\gamma\delta$T cell-specific gene[140] |
| *Ikzf3* | Correlates with $\gamma\delta$T cell functions[143] |
| *Lef1* | $\gamma\delta$T cell-specific gene[140] |
| *Egr2* | Relates to $\gamma\delta$T cell expansion[144,145] |
| *Rorc* | Required for $\gamma\delta$T cell specification and differentiation[146] |

ILC cell
| Predicted putative driver TFs | Evidence |
| --- | --- |
| *Fosl2* | Necessary for NK cell differentiation[49] |
| *Runx3* | Regulates NK cell activation[50] |
| *Meis1* | Unknown |
| *Pbx3* | Unknown |
| *Bhlhe40* | Required for NK cell normal functions[147] |
| *Gata3* | Promotes NK cell differentiation[148] |
| *Nfil3* | Essential for maintenance of NK cells[149] |
| *Jdp2* | Unknown |
| *Zscan17* | Unknown |
| *Stat4* | Induces NK cell tolerance in lung diseases[150] |

**table S3.**
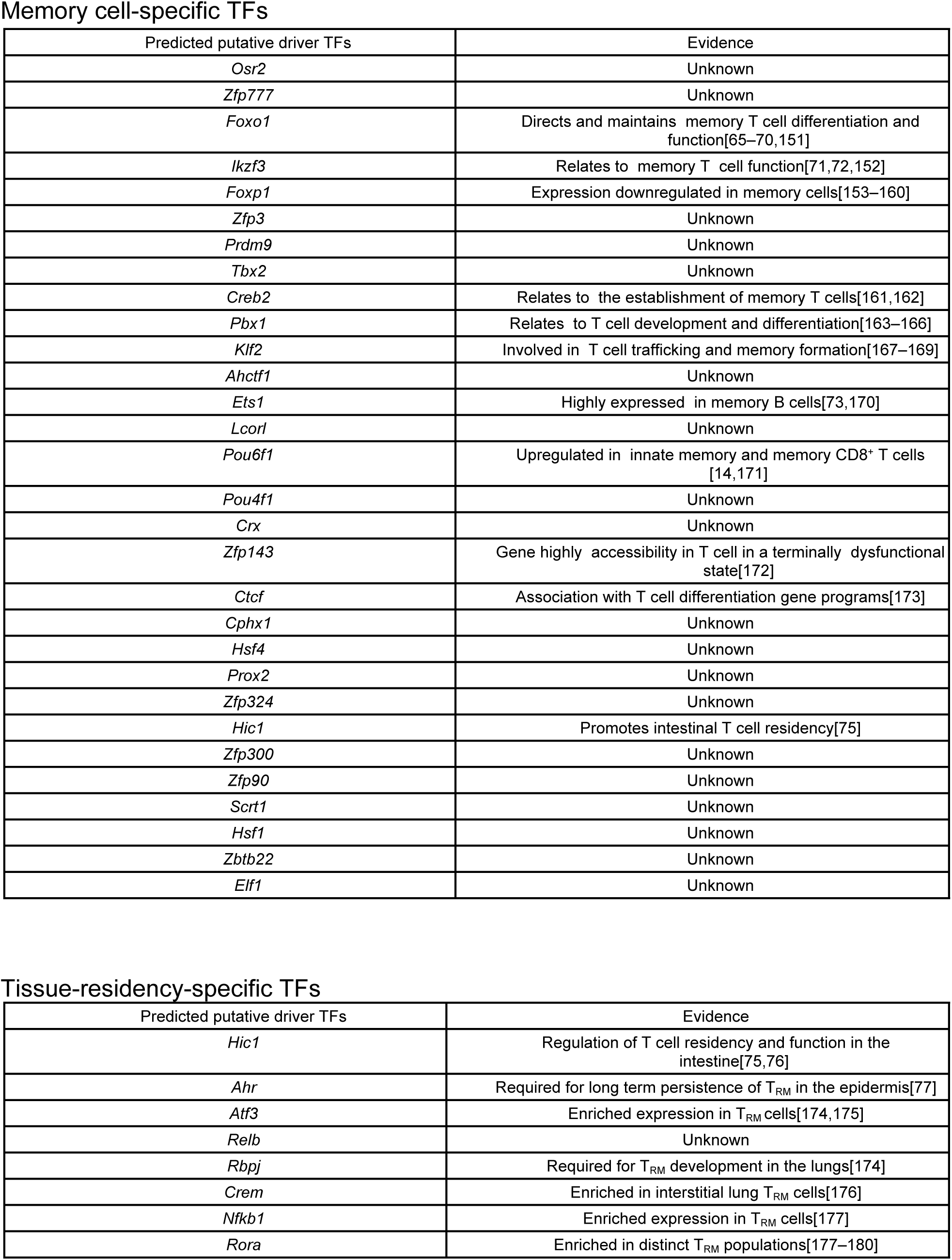
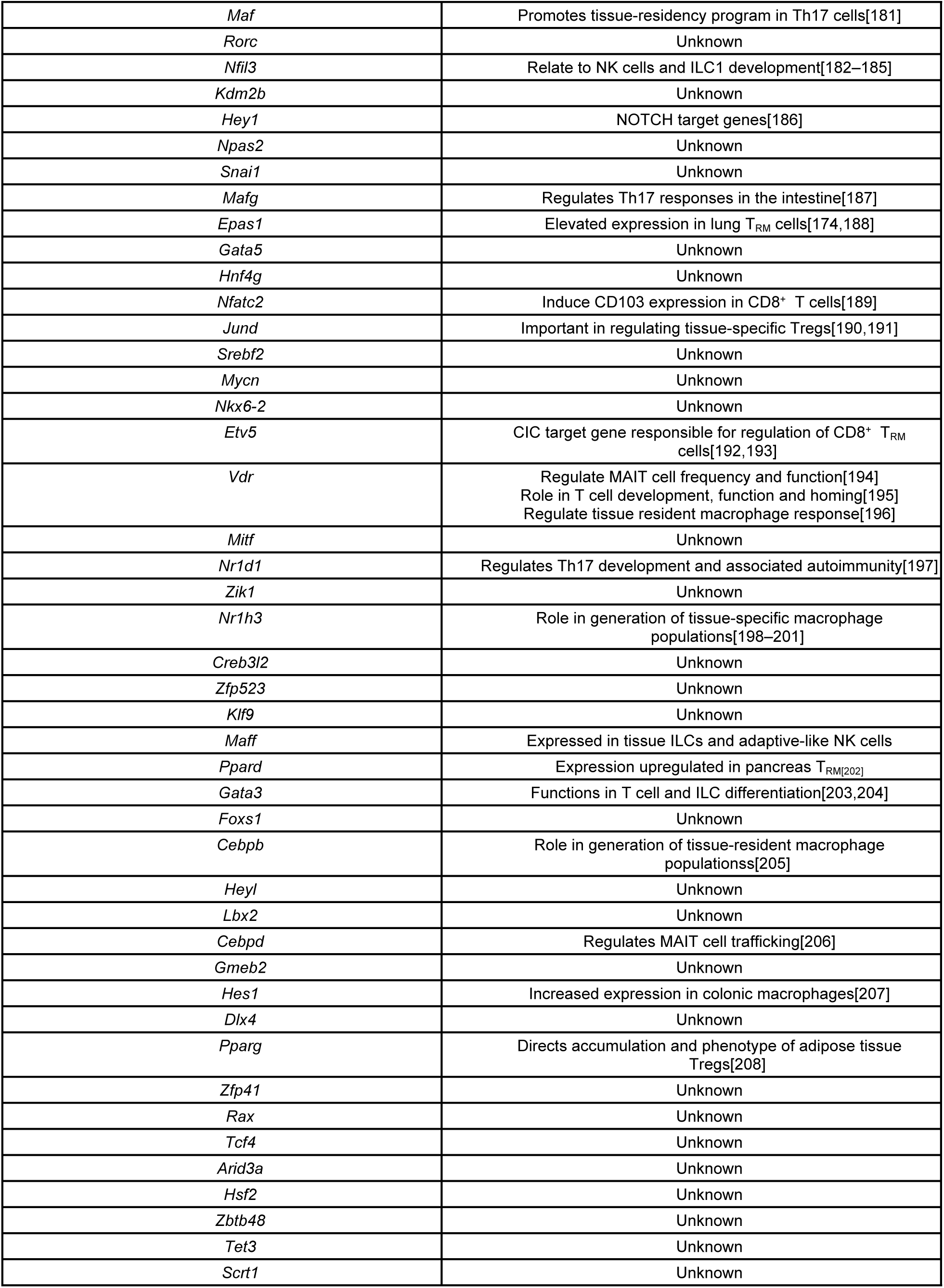
putative driver TFs in memory cells and tissue resident cells.

**table S4. GO terms and KEGG pathway enrichment statistics for the identified putative driver TFs in** **Fig. 3** **and** **Fig. 4**

[tableS4_GO_statistics.xlsx](https://docs.google.com/spreadsheets/d/1R2RJpYE0OgOcz1lzdNifdt6GlyuCiy5r/edit?usp=sharing&ouid=109046442056305381242&rtpof=true&sd=true)

